# Exploring Kp,uu,BBB Values Smaller than Unity in Remoxipride: A Physiologically-Based CNS Model Approach Highlighting Brain Metabolism in Drugs with Passive Blood-Brain Barrier Transport

**DOI:** 10.1101/2024.06.11.598191

**Authors:** Mengxu Zhang, Ilona M. Vuist, Vivi Rottschäfer, Elizabeth CM de Lange

## Abstract

**(Aim):** K_p,uu,BBB_ values are crucial indicators of drug distribution into the brain, representing the steady-state relationship between unbound concentrations in plasma and in brain extracellular fluid (brainECF). K_p,uu,BBB_ values < 1 are often interpreted as indicators of dominant active efflux transport processes at the blood-brain barrier (BBB). However, the potential impact of brain metabolism on this value is typically not addressed. In this study, we investigated the brain distribution of remoxipride, as a paradigm compound for passive BBB transport with yet unexplained brain elimination that was hypothesized to represent brain metabolism.

**(Methods):** The physiologically-based LeiCNS pharmacokinetic predictor (LeiCNS-PK model) was used to compare brain distribution of remoxipride with and without Michaelis-Menten kinetics at the BBB and/or brain cell organelle levels. To that end, multiple in-house (IV 0.7, 3.5, 4, 5.2, 7, 8, 14 and 16 mg/kg) and external (IV 4 and 8 mg/kg) rat microdialysis studies plasma and brainECF data were analysed.

**(Results):** The incorporation of active elimination through presumed brain metabolism of remoxipride in the LeiCNS-PK model significantly improved the prediction accuracy of experimentally observed brainECF profiles of this drug. The model integrated with brain metabolism in both barriers and organelles levels is named LeiCNS-PK3.5.

**(Conclusion):** For drugs with K_p,uu,BBB_ values < 1, not only the current interpretation of dominant BBB efflux transport, but also potential brain metabolism needs to be considered, especially because these may be concentration dependent. This will improve the mechanistic understanding of the processes that determine brain PK profiles.

## 1 Introduction

The challenges in accessing direct information on human brain target site concentrations have long posed a significant problem in drug development (Cook et al., 2014). To address this issue, researchers have turned to indirect approaches, utilizing methods such as observing human plasma pharmacokinetics (PK) data. However, the use of plasma PK cannot address the complex processes of blood-brain barrier (BBB) transport and intra-brain distribution (Loryan et al., 2022). To understand such PK processes, alternatively, often experimental animal data are used.

Apart from the rate of brain distribution, the extent of brain distribution, represented by the steady-state ratio of unbound concentration in the brain extracellular fluid (brainECF) over those in plasma (K_p,uu,BBB_), is a very important parameter (Hammarlund-Udenaes et al., 2008). K_p,uu,BBB_ values are often obtained from experimental animal or human cell line monolayer transport studies (Summerfield et al., 2016). When it comes to the translation and prediction of human central nervous system (CNS) compartment PK profiles, CNS physiologically-based pharmacokinetics (PBPK) models are used as a promising tool (Sanchez-Dengra et al., 2021).

For K_p,uu,BBB_ values around 1, passive BBB transport (diffusion) is the dominating process. The interpretation typically is that for K_p,uu,BBB_ values smaller than 1, active efflux transport dominates, while the other way around, for values larger than 1, active influx transport is dominant. For K_p,uu,BBB_, typically all attention has been on the impact of active transport processes. However, metabolism could also play a significant role.

More and more studies indicate the abundant presence of drug-metabolizing enzymes (DMEs) within the brain, such as cytochrome P450 (CYP) (Fanni et al., 2021) and UDP-glucuronosyltransferase (UGT) enzymes (Sakakibara et al., 2016). Emerging insights revealed that many CNS-active drugs undergo brain metabolism, like codeine (Chen et al., 1990), clozapine (Fang, 2000), alprazolam (Agarwal et al., 2008), propofol (Khokhar and Tyndale, 2011), and tramadol (Wang et al., 2015). These findings have prompted a growing recognition of the need to consider brain metabolism in conjunction with other factors when studying CNS drug distribution.

In this context, CNS PBPK model approaches have proven their added value for predicting drug brain distribution in rats, especially for their potential for translational applications to humans. Such models make the explicit distinction between drug properties and CNS physiological parameters, which are used as input variables, together with estimated plasma PK exposure that depends on dosing regimens (Saleh et al., 2021; Sanchez-Dengra et al., 2021). Notably, our group developed the most comprehensive CNS PBPK model called the LeiCNS-PK model, including 9 CNS compartments (Saleh et al., 2021; Yamamoto et al., 2018). The most advanced version, LeiCNS-PK3.0, offers comprehensive insights into drug distribution within various CNS compartments. However, so far, this model did not consider potential brain metabolism.

Remoxipride, a compound without any active transport reported, posed a challenge for PK model development of brain distribution using plasma and brain microdialysis datasets. The best-fitting models identified passive BBB transport, with the need to include brain elimination (Stevens et al., 2011; van den Brink et al., 2017a). In other words, there was a discernible loss of remoxipride from the brain that did not return to plasma (van den Brink et al., 2017b), strongly suggested the involvement of brain metabolism in the elimination process.

Remoxripride is known to be metabolized by CYP2D6 in humans (Marsh et al., 1999; Movin-Osswald et al., 1993). While concrete evidence is lacking to definitively identify the responsible rat CYP2D isoform for remoxipride metabolism (Miksys et al., 2000; Tyndale et al., 1999), it is certain that the CYP2D isoforms in rats fulfil the role of CYP2D6 in humans (Grobe et al., 2012). Although there are no studies directly measuring remoxipride metabolism in rat brain, studies on remoxipride and its brain metabolites that have revealed consistently high brain/plasma ratios for remoxipride FLA metabolites (∼10-25) (Ogren et al., 1993), strongly suggesting that localized remoxipride metabolism occurs in the brain. Furthermore, CYP2D activity in rat brain microsomes was identified using dextromethorphan O-demethylation as a probe (Jolivalt et al., 1995). Altogether, our hypothesis is that brain metabolism is responsible for brain elimination of remoxipride.

Brain expression of particular DMEs has been found to be extensive, especially for CYP enzymes (Fanni et al., 2021; Miksys and Tyndale, 2013) and UGT enzymes (Ouzzine et al., 2014; Yang et al., 2022). It has been reported that mammalian CYPs are membrane-associated enzymes, mostly attached to the endoplasmic reticula (ER) and inner membranes of mitochondria (MT) facing the cytosol space of the cell (McMillan, 2018; Šrejber et al., 2018), where the substrate binding pockets are deeply buried in the phospholipid bilayer (Šrejber et al., 2018). Therefore, we considered brain metabolism to mainly happen in the brain cell organelles’ membranes. Moreover, although limited, DMEs are also expressed within the BBB (Decleves et al., 2011) and the blood-cerebrospinal fluid barrier (BCSFB) (Wang and Zuo, 2018), where they could work as a metabolic barrier (Kadry et al., 2020).

In this study we used our LeiCNS-PK3.0 model, as the most comprehensive CNS PBPK model (Saleh et al., 2021), to compare brainECF predictions without and with metabolism, be it within the BBB/BCSFB, or within brain cell organelles’ membranes (Šrejber et al., 2018). Here we modelled metabolism by a Michaelis-Menten term. We utilized comprehensive rat brain distribution datasets obtained at different dosages to validate our model LeiCNSPK3.5. Finally, we compared the brain distribution predictions of remoxipride in brainECF by the LeiCNS-PK models, with observed brainECF PK.

## 2. Material and methods

### 2.1 Model development

#### 2.1.1 The LeiCNS-PK3.0 model

For this study the previously published CNS PBPK model, LeiCNS-PK3.0 (Figure 1A), was used as base model (Saleh et al., 2021). This comprehensive model consists of plasma, multiple CNS compartments and fluid flows between them. Specifically, these multiple physiological compartments, including brain microvasculature (MV) and several CSF compartments, are connected through cerebral blood flow, brainECF bulk flow, and CSF flow. Barriers are set according to the physiological structure, between brainMV and brainECF. Furthermore, this model considers pH values in each compartment, as well as brain non-specific tissue binding.

**Figure 1.**
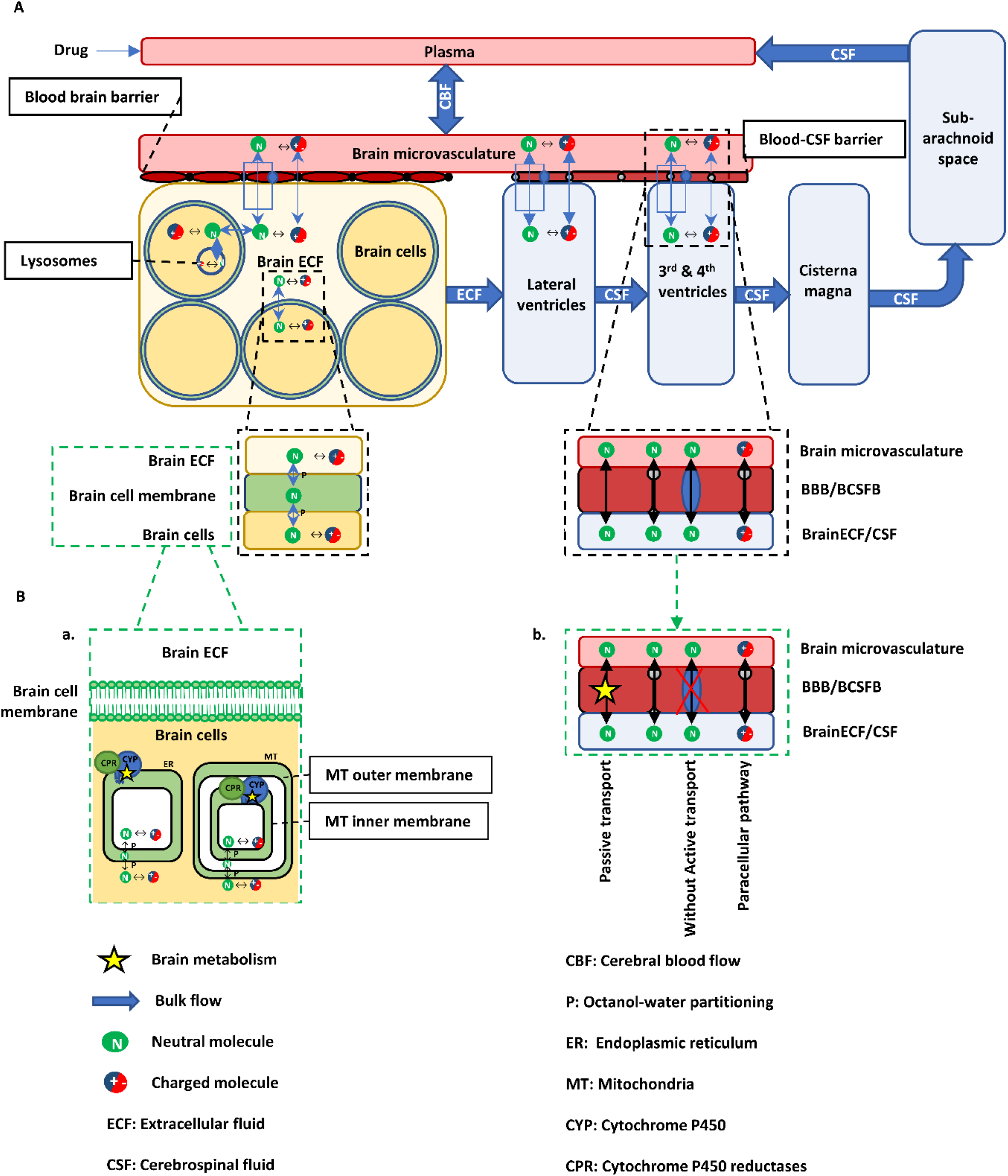
Incorporation of brain metabolism based on LeiCNS-PK3.0 model. **(A)** LeiCNS-PK3.0 is a nine compartments PBPK model, including compartments, flows in the brain, and transport mechanisms, pH factors and other physiological processes, predicting unbound drug concentrations of brain compartments in rats (Saleh et al., 2021). **(B)** We added brain metabolism for the 3.5 model by adding metabolism into newly developed compartments (a), endoplasmic reticulum (ER) and mitochondria (MT), and barriers (b) in green dashed frames. CYPs are mainly expressed in the ER and inner membrane of MT (Šrejber et al., 2018). They deeply anchored in the membrane (Durairaj and Li, 2022), and the active sites of CYP enzymes are also buried in the membrane (Šrejber et al., 2018). Thus, the metabolism that takes place in the ER and MT membranes is thought to occur when drug molecules cross the membranes. In the green dashed frame, physiological position of CYPs in brain cell organelles are presented. The detailed LeiCNS-PK3.5 model structure is shown in Figure S1.

#### 2.1.2 Incorporation of brain metabolism in the LeiCNS-PK model

To address brain metabolism in brain barriers and brain cell organelles, a Michaelis-Menten term was included (Deepika and Kumar, 2023; Lipscomb and Poet, 2008) into the LeiCNS-PK3.0 model at different locations (bright yellow stars in Figure 1B). Michaelis-Menten related parameters (V_max_ and K_m_) were considered generic, addressing overall metabolic parameters in the brain. The resulting model is named LeiCNSPK3.5

##### 2.1.2.1 Metabolism at the level of endoplasmic reticulum and mitochondria

In the LeiCNS-PK3.0 model, the brain intracellular fluid compartment (brainICF) comprises only one compartment (lysosomes). To integrate metabolism within brain cell organelles, we introduced four additional compartments within brainICF to delineate the ER, ER membrane (ERM), MT, and MT membranes (MTM).

The changes of drug amounts in each compartment (*A*_*ICF*_, *A*_*ERM*_, *A*_*ER*_, *A*_*MTM*_, *A*_*MT*_) are described by Equations 1 to 5. Here, *CL*_*WO*_ (ml/min) is the water-to-lipid clearance (flow into brain cell plasma membrane compartment, BCM), while *CL*_*OW*_ (ml/min) represents the lipid-to-water clearance (flow out of BCM). *ERCL*_*WO*_ *and MTCL*_*WO*_ indicate the water-to-lipid clearance flow into the ERM and MTM. Conversely, *ERCL*_*OW*_ *and MTCL*_*OW*_ are the lipid-to-water clearance flow out of the ERM and MTM. Since the CYP active sites are deeply embedded within the ER and inner MT membranes (Šrejber et al., 2018), drug molecules exclusively undergo metabolism within these membranes. The parameters *efSA*_*ER*_ and *efSA*_*MT*_ denote the effective metabolic surface area fraction of ERM and inner MTM, respectively.

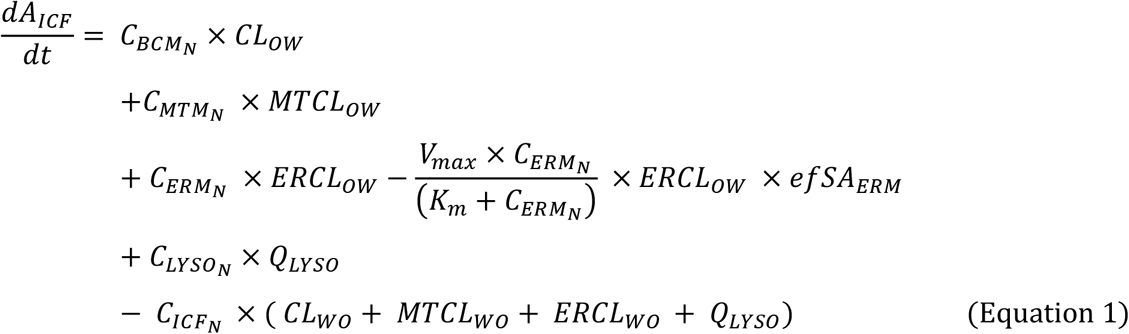

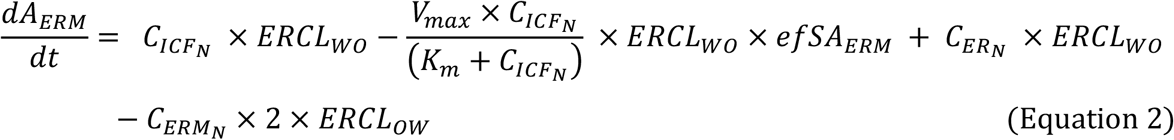

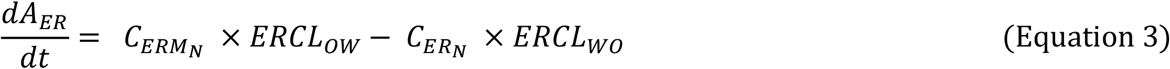

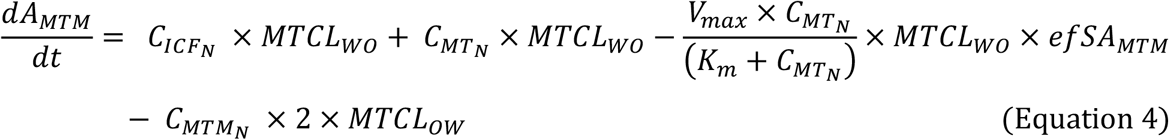

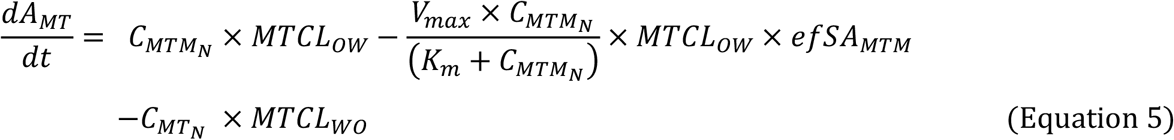

In which *V*_*max*_ (ng/ml*min) is the maximum velocity achieved by the enzyme when the drug concentration is high enough to saturate the enzyme, and *K*_*m*_ (ng/ml) is the Michaelis-Menten constant. *Q*_*LYSO*_ is the transmembrane clearance of lysosomes in ml/min. *C*_*BCM*_, *C*_*MTM*_, *C*_*ERM*_, *C*_*MT*_, *C*_*ER*_, *C*_*LYSO*_, *C*_*ICF*_ are the unbound concentrations (ng/ml) in BCM, MTM, ERM, MT, ER, lysosome (LYSO), and brainICF compartments, respectively. 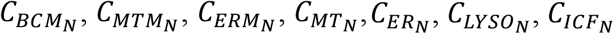 are the unbound neutral concentrations in corresponding compartment.

##### 2.1.2.2 Metabolism at the level of BBB and BCSFB

In the LeiCNS-PK3.0 model, BBB and BCSFB transport is incorporated using asymmetry factors (AFs). To include metabolism at the level of the BBB/BCSFB we need to reconsider the AFs. After all existing physiological processes in the model are excluded, an AF value remains as the *“pure”* extent of drug distribution at a particular barrier. In the LeiCNS-PK3.0 model, the AFs are calculated using K_p,uu,BBB_ values with explained physiological processes as input (see Figure 2). But, as indicated earlier, for K_p,uu,BBB_ values lower than one, brain metabolism is typically not considered, and should be explicitly addressed. Therefore, we revised the AFs, by taking metabolism out of the AFs, with a separated Michaelis-Menten term representing metabolism. Thus, in the new model (LeiCNSPK 3.5), the AFs exclusively encompasses factors related to active transport across barriers and any other as-yet-unknown physiological processes that may affect drug concentrations in the brain. In this context, we re-evaluate and determine a new constant AF.

**Figure 2.**
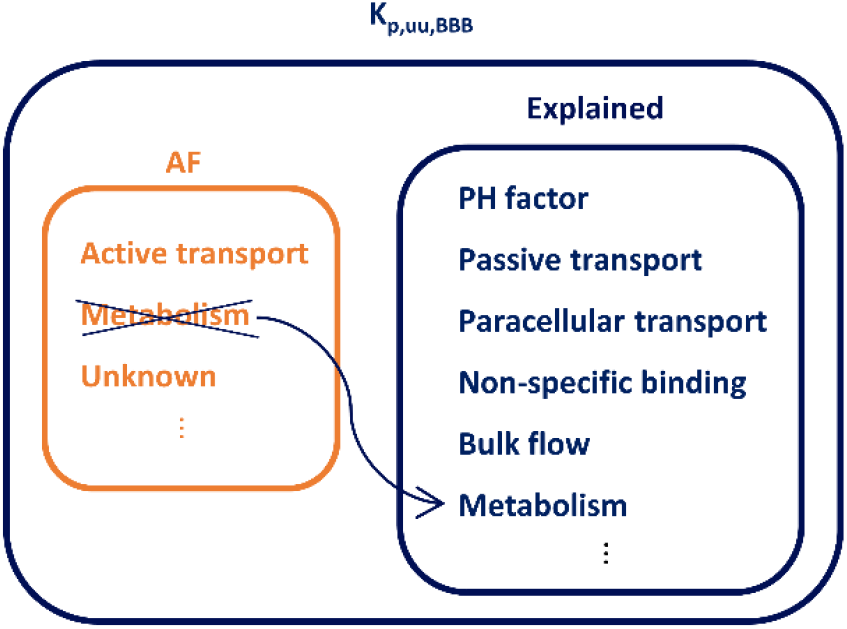
The relationship between K_p,uu,BBB_ and all other physiological parameters under the *in vivo* steady-state, including AF. The orange colour represents the temporarily unexplained physiological processes. The dark blue colour represents the explained physiological processes in the LeiCNS-PK3.0 model. The difference between the LeiCNS-PK3.0 and the LeiCNS-PK3.5 is brain metabolism. To that end, metabolism is taken out from AF and incorporated it in the “explained” part. Thereby, brain metabolism is an explicitly defined physiological process in the PBPK model.

As mentioned before, according to previous PK modelling of microdialysis data remoxipride was identified to cross the BBB passively (Stevens et al., 2011; van den Brink et al., 2017a). Therefore, we set AFs (*AF*_*in*_ and *AF*_*ef*_) values equal to 1, which means there is no active transport at the level of the brain barriers.

Previously, the amount per time of drug transcellularly crossing the BBB and BCSFB was modelled by *Q*_*t*_ × *AF* × *C*_*N*_, where *C*_*N*_ is the unbound neutral concentration (ng/ml) of drug. In equation 6, the changes of the drug amount in brainMV compartment *A*_*MV*_ (in ng) is described. Equation 7 quantifies the amount per time of drug in the brainECF *A*_*ECF*_ (in ng). *Q*_*t*_ is the transcellular transport clearance in ml/min, encompassing both passive and active transcellular components. *Q*_*p*_ is the paracellular transport clearance in ml/min. *AF*_*in*_ is the influx asymmetry factor as mentioned before. *AF*_*ef*_ is the efflux asymmetry factor.

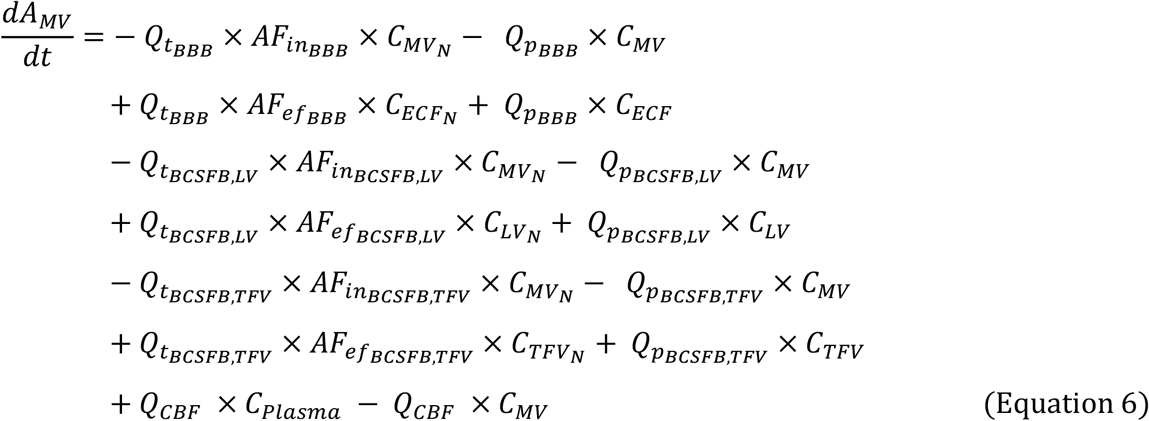

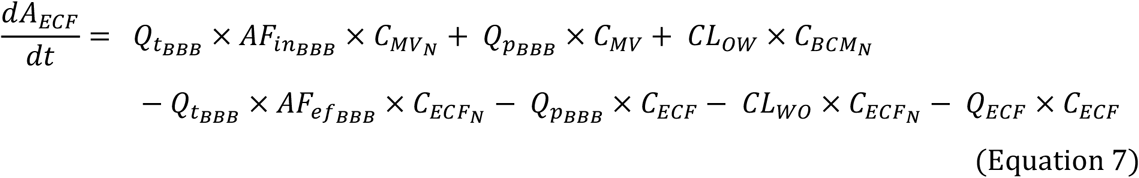

In the new model, we added brain metabolism within the barriers. The elimination of drug molecules by brain metabolism is quantified by including a Michaelis-Menten term into equations 6 and 7, resulting in equations 8 and 9.

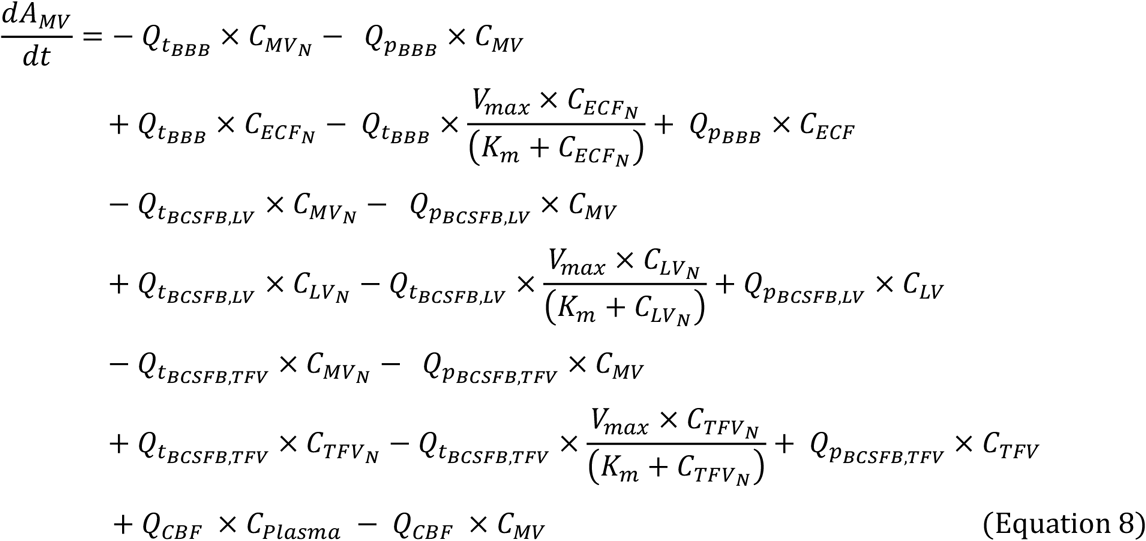

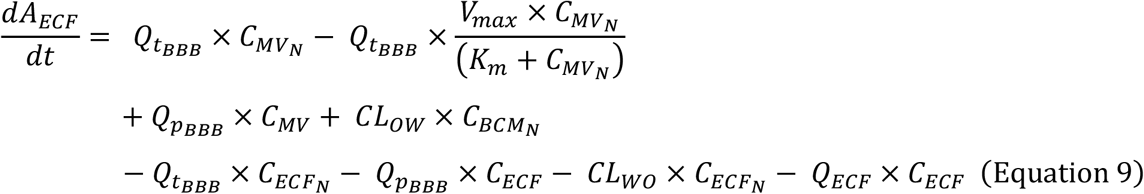

In equations 8 and 9, there are no AF values present because, for remoxipride, as mentioned above, these are set equal to 1. *Q*_*t*_ is still the transcellular clearance, but only the passive transcellular component is left. *C*_*MV*_, *C*_*ECF*_, *C*_*LV*_, *C*_*TFV*_ and *C*_*plasma*_ are the unbound concentrations of the drug in the brainMV, brainECF, brain lateral ventricles (brainLV) and brain 3^rd^ and 4^th^ ventricles (brainTFV), and plasma compartments, respectively. 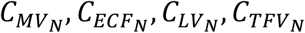are the unbound concentrations of the neutral drug in the corresponding compartment. After the model development in which we added metabolism in organelles’ and barriers’ levels to the LeiCNS-PK3.0 model, we name the new model **LeiCNS-PK3.5**.

### 2.2 LeiCNS-PK model input

In general, the input data for the LeiCNS-PK model is divided into three parts: drug-specific parameters (Table S1), system-specific parameters (Table S2), and plasma PK parameters (in section 4.1). Specifically for the LeiCNSPK3.5, remoxipride Michaelis-Menten parameters were added. All other drug-specific and system-specific parameters come from the Drugbank database (Wishart et al., 2018) and literature, respectively.

#### 2.2.1 Remoxipride Michaelis-Menten parameters

We constructed a 3-compartment brain metabolism model (Figure S2 and Figure S3) employing non-linear mixed effects modelling to derive the brain’s K_m_ and V_max_ values (Table S3) based on observed rat plasma and microdialysis brainECF PK data from 4 in-house studies (Table 1). The software is mentioned in the “Data analysis and software” section.

**Table 1.**
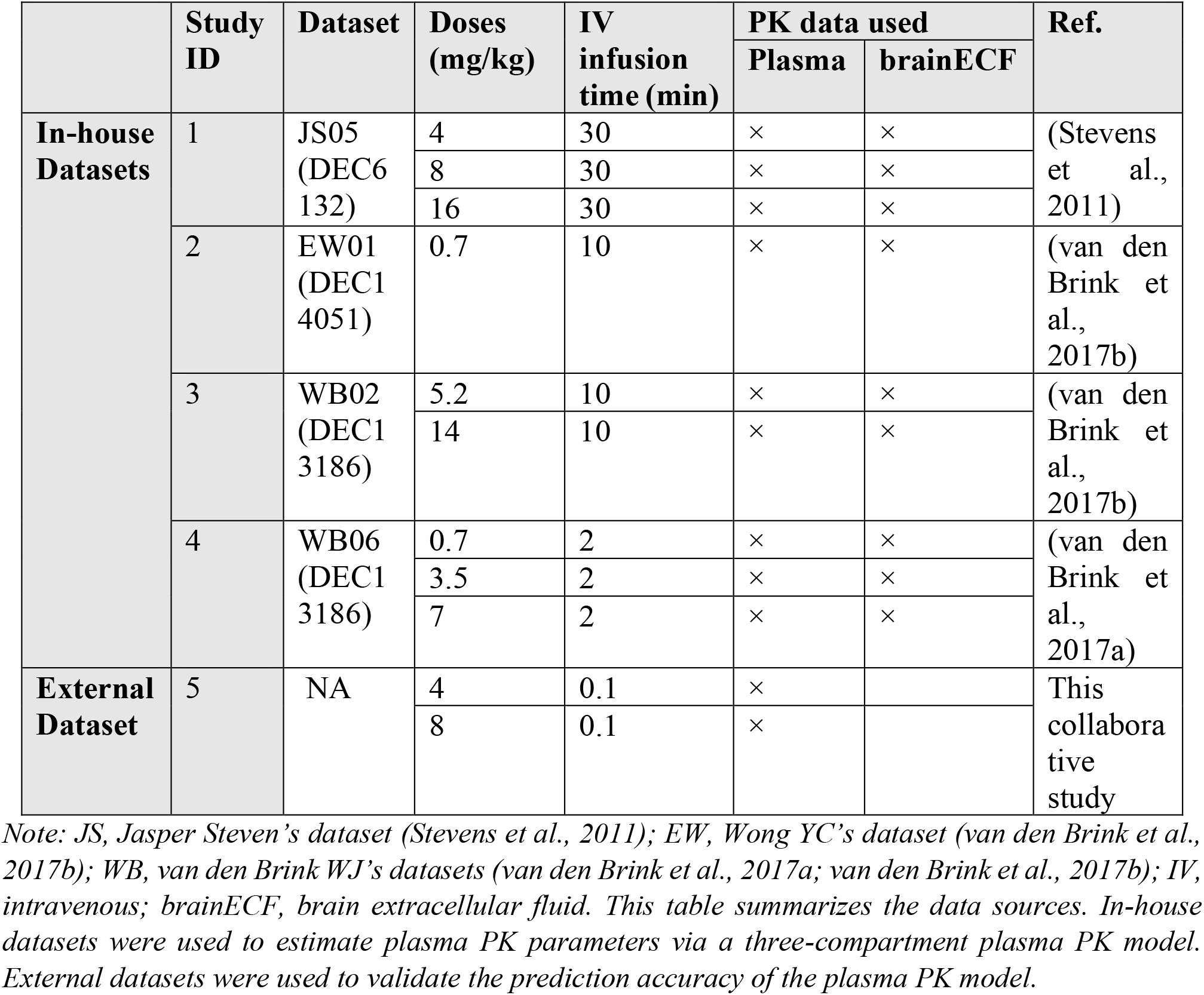
Summarization of remoxipride plasma PK and brainECF PK rat datasets used in this project.

|  | Study ID | Dataset | Doses (mg/kg) | IV infusion time (min) | PK data used |  | Ref. |
| --- | --- | --- | --- | --- | --- | --- | --- |
|  |  |  |  |  | Plasma | brainECF |  |
| In-house Datasets | 1 | JS05 (DEC6 132) | 4 | 30 | × | × | (Stevens et al., 2011) |
|  |  |  | 8 | 30 | × | × |  |
|  |  |  | 16 | 30 | × | × |  |
|  | 2 | EW01 (DEC1 4051) | 0.7 | 10 | × | × | (van den Brink et al., 2017b) |
|  | 3 | WB02 (DEC1 3186) | 5.2 | 10 | × | × | (van den Brink et al., 2017b) |
|  |  |  | 14 | 10 | × | × |  |
|  | 4 | WB06 (DEC1 3186) | 0.7 | 2 | × | × | (van den Brink et al., 2017a) |
|  |  |  | 3.5 | 2 | × | × |  |
|  |  |  | 7 | 2 | × | × |  |
| External Dataset | 5 | NA | 4 | 0.1 | × |  | This collaborative study |
|  |  |  | 8 | 0.1 | × |  |  |

*Note: JS, Jasper Steven's dataset (Stevens et al., 2011); EW, Wong YC's dataset (van den Brink et al., 2017b); WB, van den Brink WJ's datasets (van den Brink et al., 2017a; van den Brink et al., 2017b); IV, intravenous; brainECF, brain extracellular fluid. This table summarizes the data sources. In-house datasets were used to estimate plasma PK parameters via a three-compartment plasma PK model. External datasets were used to validate the prediction accuracy of the plasma PK model.*

#### 2.2.2 Plasma PK parameters’ estimation

##### 2.2.2.1 Data for remoxipride plasma PK model

Our research benefits from 4 previous in-house studies conducted by colleagues within our research group. Studies 1 to 4 (listed in Table 1) have produced observed rat plasma PK data regarding remoxipride. We utilized these data to develop a plasma PK model, from which critical plasma PK parameters were derived to serve as input for the LeiCNS-PK3.0 and 3.5 models. Study 5 was an externally performed and those data were used as an external dataset to validate the estimation of plasma PK parameters determined on the internal datasets.

##### 2.2.2.2 Plasma PK modelling

The rat plasma PK model was constructed, using non-linear mixed effects modelling, to provide input for the LeiCNS-PK models. The suitability of 1-, 2-, and 3-compartment models was compared to identify the best-fitting model for the data. Finally, the three-compartment model was selected, see Figure 3. An exponential method was employed, enabling the incorporation of individual differences in drug response. Furthermore, to account for unexplained variability and random errors within the data, a proportional error model was implemented.

**Figure 3.**
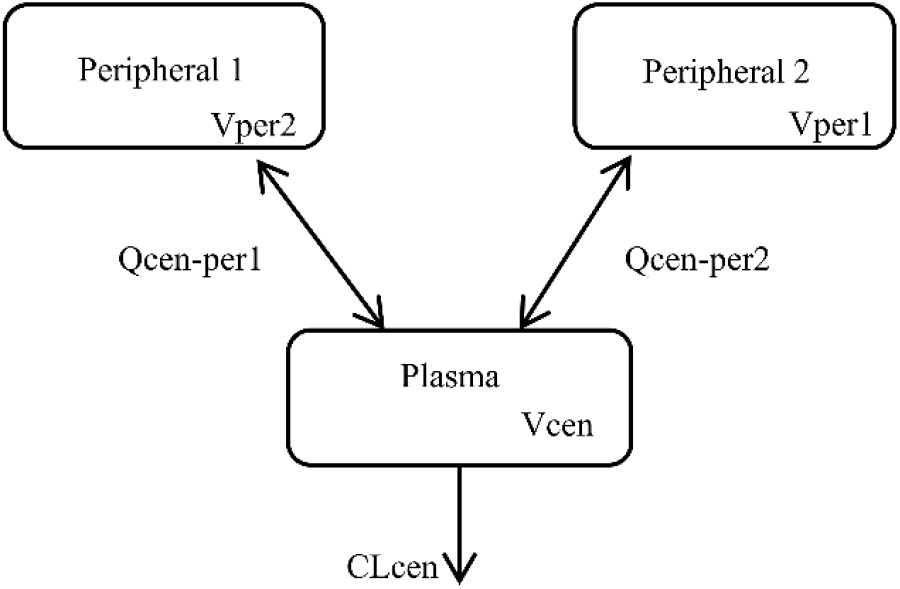
The empirical plasma PK model structure. A 3-compartmental plasma PK model was chosen according to the model evaluation process described above. Clcen, clearance from central compartment; Qcen-per1, clearance between central compartment and peripheral 1 compartment; Qcen-per2, clearance between central compartment and peripheral 2 compartment; Vcen, distribution volume in central compartment; Vper1, distribution volume in peripheral 1 compartment; Vper2, distribution volume in peripheral 2 compartment.

The final selection of the plasma PK model was based on a comprehensive set of criteria. A likelihood ratio test with a significance level of p < 0.05, along with a significant decrease in the Akaike information criterion (AIC), was used to determine if the more complex model (3-compartment model) provided a significantly better fit compared to a simpler alternative (one- or two-compartment model). Additionally, visual predictive check (VPC) plots were utilized to evaluate the model’s performance in predicting drug concentrations in plasma. The precision of parameter estimates was denoted by % relative standard errors (%RSE), with lower values indicating more precise estimates. To further assess the model’s accuracy and goodness of fit, various plots were generated (Figure S4A-S4C), including observations versus individual and population predictions, and population weighted residuals versus population prediction over time. Other pertinent factors such as correlations, condition number and shrinkage were also taken into consideration during the model selection process. Upon model development, the selected 3-compartment plasma PK model successfully described remoxipride plasma concentrations as input for the LeiCNS-PK models.

### 2.3 External dataset materials and methods

The external dataset (study 5) was generated specifically for this collaborative study. Remoxipride PK profiles were measured under 2 different dosages (4 mg/kg and 8 mg/kg) in rats, providing plasma and brainECF data at multiple time points. Details on experiments of this dataset are in the supplementary file “*Experimental Details*” section.

### 2.4 The LeiCNS-PK model accuracy evaluation, Comparison between 3.0 and 3.5

We employed the above input data and experimental observed PK data to assess the predictive performance of the LeiCNS-PK3.5, in which brain metabolism is included. This performance was compared with that of the LeiCNS-PK3.0 version. We achieved this evaluation by quantifying the disparities between the model’s predictions and the actual observations.

The LeiCNS-PK3.5 model’s performance evaluation involved comprehensive statistical analysis, assessing accuracy, bias, and variability in predicting drug concentrations. Key statistical methods ensured thorough examination. VPC plots played a pivotal role. These plots visually represented model prediction alignment by comparing the median and 95% prediction interval from 200 simulations with *in vivo*-measured unbound drug concentrations, offering an initial performance overview. To handle interindividual variability (η) and residual variabilities (ε) in the plasma PK model, simulations utilized random sampling from normal distributions with mean of 0 and variances of ω2 and σ2, respectively. This ensured proper consideration of individual differences and unexplained variations within the plasma PK model.

Prediction errors were computed by comparing individual measured drug concentrations with corresponding time-matched simulation medians. The relative accuracy (RA_drug_), computed for each dose and compartment, averaged logarithmic differences between all observed drug concentrations and simulated median concentrations. RA_drug_ values nearing 0 affirmed accurate, reliable predictions, reinforcing model validity. Average fold error (AFE) and absolute average fold error (AAFE) were derived from RA_drug_. AFE, representing relative accuracy for all dosages, quantified model bias, with values near 100% indicating minimal bias. AAFE gauged how well simulated typical PK profiles matched measured typical PK data, with values close to 100% indicating accuracy. These summary metrics offered a concise assessment of model accuracy, facilitating comparisons across dosages.

RA_drug_ at a given compartment was calculated as follows:

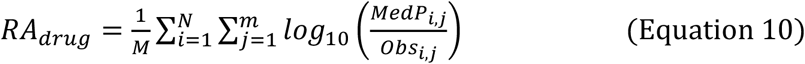

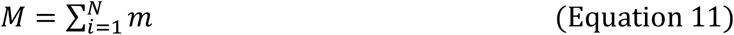

Here Obs_i,j_ (or MedP_i,j_) represents j^th^ observation (or median value of 200 simulations corresponding to Obs_i,j_) of the i^th^ individual; M stands for the total number of observations across all individuals; N indicates the overall number of individuals; and m signifies the count of observations of the i^th^ individual.

AFE of specific compartment was calculated as:

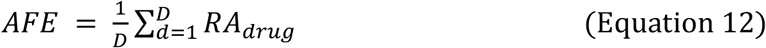

Here D is the number of dosages used for evaluation.

AAFE of a specific compartment was calculated as:

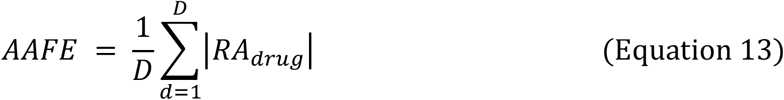

### 2.5 Data analysis and software

The non-linear mixed effect model were built up using Monolix and Sycomore version 2021R2. For general data analysis, visualization, LeiCNS-PK3.0 simulations, and LeiCNS-PK3.5 simulations, R version 4.3.0 (R Core Team, 2023) was employed. The simulations were carried out using the rxode2 package version 2.0.13 (Wang et al., 2016), and the LSODA (Livermore Solver for Ordinary Differential Equations) Fortran package was utilized for solving the differential equations. Furthermore, algebraic equations were solved using Maxima Computer Algebra System version 22.04.0 (available from http://maxima.sourceforge.net).

## 3 Results

### 3.1 Plasma PK parameters

The rat plasma PK model parameters were estimated with high precision and an accurate description of observed plasma drug concentrations (Table 2).

**Table 2.** Rat plasma PK parameters used as input to LeiCNS-PK model.

| REMOXIPRIDE | PARAMETERS | VALUES | STOCH.APPROX. |  |
| --- | --- | --- | --- | --- |
|  |  |  | S.E. | R.S.E. (%) |
| Estimated Plasma PK Typical Values | Clcen (mL/min) | 32.46 | 1.25 | 3.84 |
|  | Qcen-per1 (mL/min) | 11.01 | 0.31 | 2.80 |
|  | Qcen-per2 (mL/min) | 50.73 | 3.47 | 6.85 |
|  | Vcen (mL) | 1.64 | 0.21 | 13.0 |
|  | Vper1 (mL) | 405.06 | 17.2 | 4.25 |
|  | Vper2 (mL) | 338.07 | 22.44 | 6.64 |
| **Standard Deviation of the Interindividual Variability** | **Omega_Clcen (mL/min)** | 0.34 | 0.028 | 8.2 |
|  | **Omega_Qcen-per1 (mL/min)** | 0.08 | 0.022 | 27.1 |
|  | **Omega_Qcen-per2 (mL/min)** | 0.37 | 0.073 | 20.0 |
|  | **Omega_Vcen (mL)** | 0.52 | 0.11 | 21.3 |
|  | **Omega_Vper1 (mL)** | 0.3 | 0.035 | 11.7 |
|  | **Omega_Vper2 (mL)** | 0.47 | 0.054 | 11.5 |
| **Residual Variability** | **Proportional** | 0.19 | 0.0077 | 3.99 |

*Note: Parameters are corresponding to Figure 3. The proportional residual variability was chosen to describe the unexplained differences between the observations and the individual predictions (Proportional). STOCH.APPROX, stochastic approximation; S.E., standard error; R.S.E, relative standard error. The parameters were estimated using the basic 3-compartmental model in Figure 3.*

### 3.2 LeiCNS-PK model predictions – comparison without (version 3.0) and with (version 3.5) Michaelis-Menten kinetics

Before predicting the brainECF remoxipride concentrations using the LeiCNS-PK3.5 model, we validated our estimated plasma PK parameters for being appropriate as input. We can see in the Figure 4A, that our estimated plasma PK parameters by internal datasets (study 1 to 4) can properly describe the measured remoxipride concentrations in plasma over time in both 4 mg/kg and 8 mg/kg dosages (study 5).

**Figure 4.**
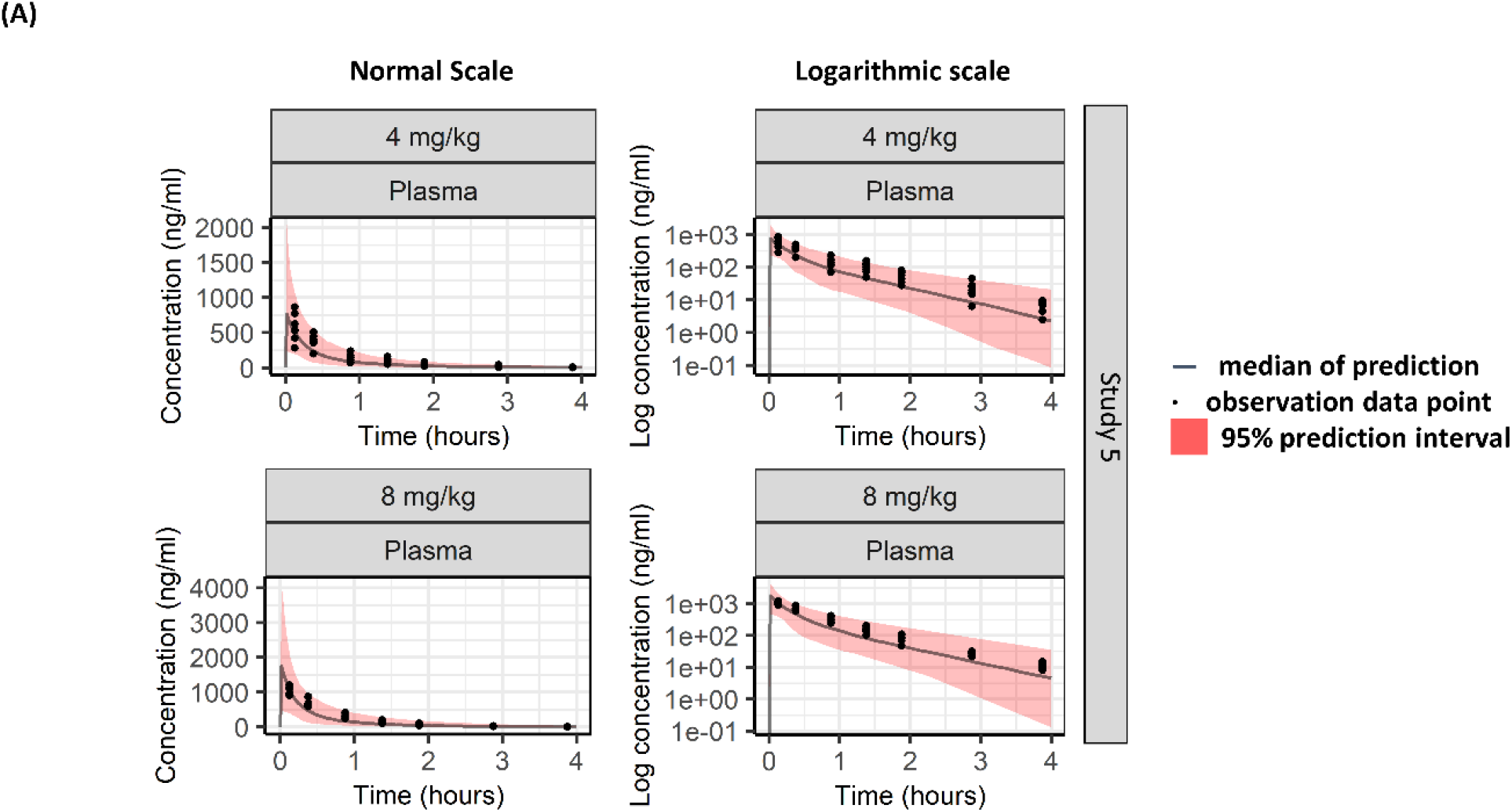
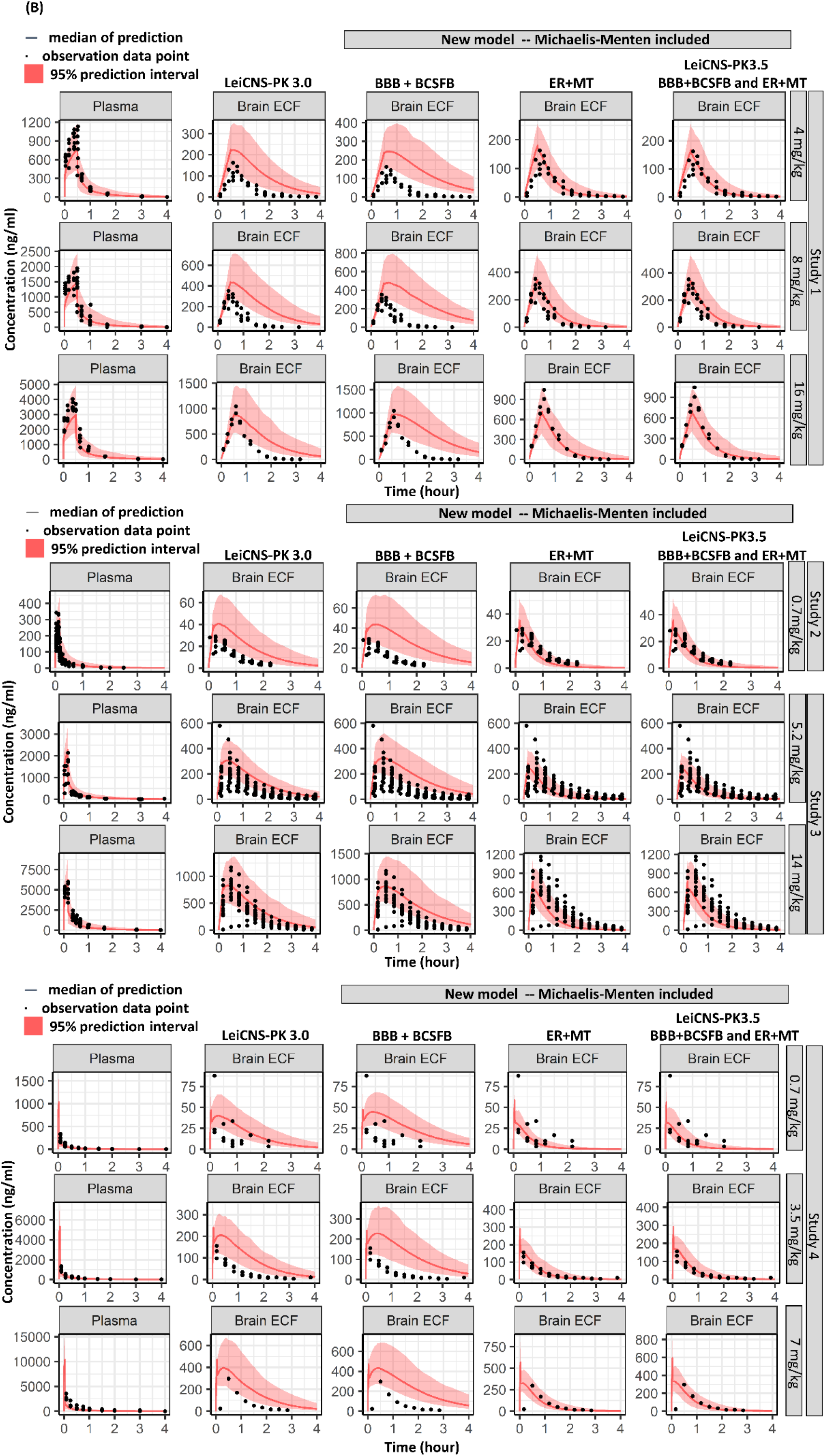
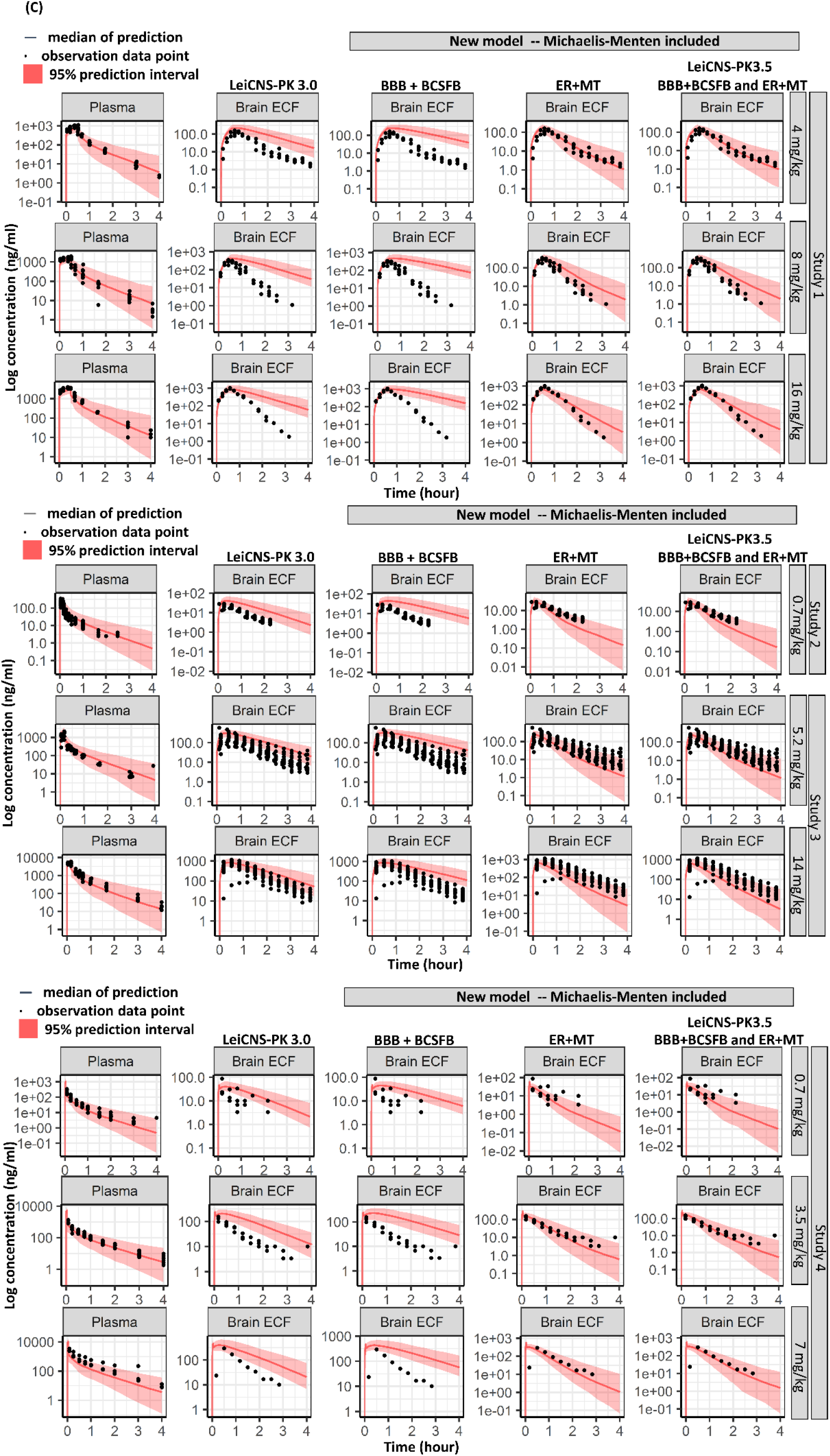
Validation of estimated plasma PK parameters and comparisons of remoxipride brainECF concentration predictions between LeiCNS-PK3.0 (without metabolism) and LeiCNS-PK3.5 model (with metabolism). (A) Estimated plasma PK parameters are validated using an external dataset via visual prediction check (VPC). The external dataset is not included in building up and estimating remoxipride plasma PK parameters. Besides, to better understand metabolic processes, we performed a VPC by comparing the LeiCNS-PK model without and with metabolism (3.0 vs. 3.5 model) on a normal scale (B) and a logarithmic scale (C). Predictive accuracy is shown by comparing in vivo measured drug concentrations in plasma (A) and brainECF (B, C) (indicated by black dots) with the median (solid line) and 95% prediction intervals (red bands) of 200 model simulations. This analysis (B and C) spanned four studies and nine different doses of remoxipride. The VPC plots arrange columns as follows: plasma data, LeiCNS-PK3.0 predictions, metabolism only in barriers, metabolism solely in ER and MT, and metabolism within ER, MT, and barriers (LeiCNS-PK3.5), offering a comprehensive view of major metabolic sites.

After preparing all the input data, predictions of brainECF remoxipride concentrations using LeiCNS-PK models (3.0 and 3.5) were executed. The plasma and brainECF predictions of remoxipride in LeiCNS-PK (3.0 versus 3.5) are presented in Figures 4B and 4C. It can be seen that the LeiCNS-PK3.5 shows a substantially improved prediction of the observed data in the brainECF for all dosages of remoxipride in brainECF compartment compared to LeiCNSPK3.0.

Also, it can be seen that, for remoxipride, metabolism in the brain barriers does not play a significant role, leading to very similar results between the Michaelis-Menten added in “BBB+BCSFB” and LeiCNS-PK3.0 simulations, and the Michaelis-Menten added in “ER+MT” and “BBB+BCSFB and ER+MT” in Figure 4B-4C. Thus, based on the results in Figure 4, combined with what is already known, we conclude that, for remoxipride, the primary metabolic location in the brain is at the level of the brain cells on the ER and MT, not in the barriers.

### 3.3 Rat LeiCNS-PK model evaluation

In addition, the LeiCNS-PK3.5 model’s performance was evaluated by calculating the RA_drug_ error, AFE and AAFE. AFE and AAFE assess the model’s bias and typical PK profile predictability, respectively (Figure 5).

**Figure 5.**
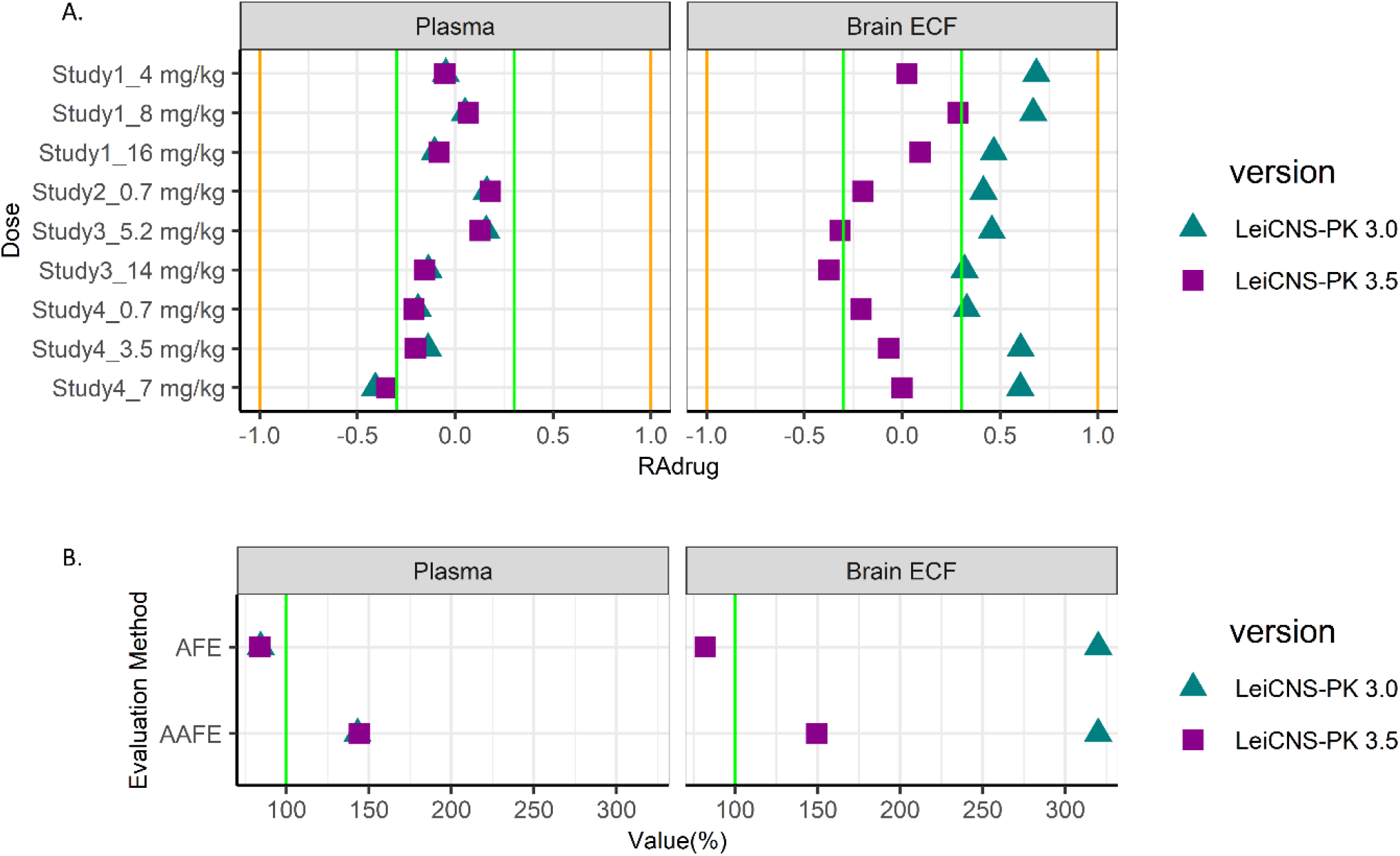
The RA_drug_, AFE and AAFE evaluations of remoxipride between LeiCNS-PK3.0 (without metabolism) vs 3.5 (with metabolism). The comparison of models using (A) RA_drug_, (B) AFE and AAFE evaluations. Green lines in A means the 2-fold error edge. The orange lines in A mean 10-fold errors. In B, values close to 100% (green line) indicated accuracy.

Here we observe that the LeiCNSPK3.5 prediction of remoxipride concentrations in brainECF presents a better accuracy and smaller bias than those of the LeiCNS-PK3.0 model. However, the RA_drug_ errors of 14 mg/kg remoxipride brainECF compartments’ predictions are slightly over the 2-fold error interval. Overall, the LeiCNSPK3.5 model predictions of other dosages still perform better than the LeiCNSPK3.0 model. All the AFE and AAFE errors in the prediction of brainECF concentrations by LeiCNSPK3.5 are substantially reduced compared with LeiCNSPK3.0.

## 4 Discussion

While the existence of DMEs in the brain is widely acknowledged (Fanni et al., 2021; McMillan and Tyndale, 2018; Miksys and Tyndale, 2002; Ouzzine et al., 2014; Sakakibara et al., 2016), drug metabolism within the brain has not received adequate attention. In this study we investigated the potential impact of brain metabolism on brainECF concentrations, and therewith on K_p,uu,BBB_. We choose remoxipride as the paradigm drug for its passive BBB and BCSFB transport and substantial brain elimination, as found in earlier studies. It was found that inclusion of active elimination of remoxipride in the brain, by readdressing AF and using a Michaelis-Menten term in the LeiCNS-PK predictor, substantially improved the prediction of observed brainECF profiles of many in-house and external datasets.

For this study we rewrote parts of the LeiCNS-PK3.0 to include the specific mathematical equations representing physiological processes of brain metabolism. In the LeiCNS-PK3.5, we explicitly separated brain metabolism from the AF. This redefines the AF, where metabolism is no longer lumped into this value, thereby primarily reflecting the remaining impact of active transport, which could be neglected for remoxipride. Consequently, the 3.5 model version considers brain metabolism when predicting unbound drug concentration in the brainECF, using a Michaelis-Menten term.

The LeiCNS-PK3.5 adequately predicts the observed brainECF data of remoxipride as obtained from multiple experimental studies, despite one outlier in RA_drug_ values for the 14 mg/kg dose. Therefore, it seems that brain enzyme conversion of remoxipride significantly affects remoxipride brainECF concentrations. Yet, the possibility of remoxipride efflux from the brain via an unidentified transporter at the BBB cannot be excluded. However, as mentioned before the mechanism-based models that investigated such a possibility did not find a basis for that (Stevens et al., 2011; van den Brink et al., 2017a).

Besides, the underprediction observed in plasma PK predictions in study 1 at 4 mg/kg and 16 mg/kg doses was attributed to using typical plasma PK parameter values instead of individual values. Figure S4D shows that individual predictions accurately fit the observed plasma data for each rat at these doses, while population predictions show underpredictions. This indicates that although the distribution of individual random effects (ŋ) is normally distributed with a mean of zero, there is a shift in ŋ within the 4 mg/kg and 16 mg/kg subgroups, particularly in CLcen and Vper1 (Figure S4E). Initially, estimating parameters for this subgroup separately was considered, but the ŋ shift issues persisted, possibly due to data imbalance from the limited sample size. Moreover, this approach did not align with our objective of uniformly describing remoxipride plasma PK parameters. Consequently, we tried covariate analysis. Preliminary assessments suggested the underprediction might result from overestimating Vcen. Although the original suspicion was dose dependency, no discernible trend in the Vcen against dosage was observed. Subsequently, the variation in rat types—Wistar WU rats in study 1 versus Wistar rats in other studies—was considered a potential factor. Attempts to incorporate rat types as a categorical covariate on Vcen did not substantially improve prediction accuracy and increased the AIC value. Further analyses introduced inter-occasion variability (IOV, different studies) or body weight as a covariate on CLcen or V3, respectively, according to Figure S4E. However, neither addition addressed the underprediction issue, and even gave unreliable parameters (high RSE%) or unstable models (condition numbers exceeding 1000). Therefore, the basic model estimated parameters (Table 2) were retained as input for the LeiCNS-PK models. Considering the accurate predictions for external data (study 5), interindividual variability, insufficient sample size of the subgroup, or lack of subpopulation-specific covariates might be the primary cause of the underprediction observed in study 1 at 4 mg/kg and 16 mg/kg doses.

It is worth noting that the LeiCNS-PK3.5 model developed is only suitable for CYP-mediated brain-metabolized drugs. If the drug is mainly metabolized by other enzymes like UGTs, the model needs to be adapted because the location of UGT enzymes is different from that of CYP enzymes.

An important limitation is the collection of the K_m_ and V_max_ parameter values. Metabolic kinetic parameters from brain are rare. Here, these parameters have been estimated by a 3-compartment PK model fit (Figure S2), and represent lumped enzymatic activity, without specific enzyme identity nor CNS anatomical locations. Therefore, these values cannot yet be considered as physiology-based. Still, combining the PBPK model with estimated parameters can accurately predict drug brainECF PK profiles and underscores the need to consider brain metabolism in CNS, and rethinking interpretation mechanisms underlying the value of K_p,uu,BBB_ or K_p,uu,brain_ values.

Comparing model predictions on brainECF PK, Michaelis-Menten kinetics only at ER and MT, or at BBB/BCSFB alone, and their combinations, indicated that for remoxipride metabolism at the level of organelles in brain cells is most important. However, it should be noted, that the inclusion of the enzyme kinetics at the level of the BBB/BCSFB, at this stage, is not defined at the level of the membranes of these barrier cells’ subcellular compartments. This is since BBB and BCSFB are not defined as cellular compartments in the LeiCNS-PK3.0 model. This may not be much of an issue as it has been reported that CYP enzyme expression in BBB/BCSFB are rare (Decleves et al., 2011; Wang and Zuo, 2018), and may not be important for remoxipride, but it might be for other drugs.

An interesting future approach would be to use the LeiCNS-PK3.5 model for the exploration of potential brain metabolism of other passively transported drugs for which K_p,uu,BBB_ values are reported as being lower than 1, provided that typically available enzymatic kinetic parameters can be found, or using brainECF data when available to be subjected to mathematical modelling as was done here for remoxipride. Furthermore, the LeiCNS-PK3.5 model provides the possibility to explore the influence of enzyme polymorphism-related individual genetic variations. For example, CYP2D6 is the major metabolic enzyme for a number of CNS active drugs (Ayano, 2016) that displays genetic polymorphism (van der Lee et al., 2021) leading to individual variations in the capability of brain metabolism. Scaling the metabolic parameters (K_m_ and V_max_) in the model according to the individual’s phenotypes or genotypes could help to predict the unbound drug PK changes in the brain for individuals.

## 5 Conclusions

For drugs with K_p,uu,BBB_ values < 1, not only the current interpretation of dominant BBB efflux transport, but also potential brain metabolism needs to be considered, especially because these may be concentration dependent. In conclusion, this study underscores the importance of considering brain metabolism in CNS modelling and simulations of drug transport into and within the brain, as well as a modelling approach how to understand its impact on brain PK. This will undoubtedly contribute to more accurate predictions and better insights into CNS drug exposure.

## 6 Acknowledgements

The first author sincerely thanks the China Scholarship Council (CSC) for providing financial support for her PhD.

## SUPPLEMENTARY MATERIALS

**Experiments Details:** MATERIAL AND METHODS FOR EXPERIMENTAL ASSESSMENT OF REMOXIRPIDE

**Table S1**. Physio-chemical parameters of remoxipride.

**Table S2**. LeiCNS-PK physiological parameters values of rats.

**Figure S1**. Detailed mathematical structure of LeiCNS-PK3.0 and the addition with brain metabolism

**Figure S2**. 3-compartment model for estimating brain metabolism kinetic parameters

**Table S3**. Parameters of 3-compartment model for remoxipride brain metabolism.

**Figure S3**. Model evaluation plots of the 3-compartment brain metabolism model

**Figure S4**. Model evaluation plots of the rat plasma PK model

## REFERENCES

Monolix 2021 R2. Lixoft SAS, a Simulations Plus company.

Sycomore 2021 R2. Lixoft SAS, a Simulations Plus company.

Agarwal, V., Kommaddi, R.P., Valli, K., Ryder, D., Hyde, T.M., Kleinman, J.E., Strobel, H.W., Ravindranath, V., 2008. Drug metabolism in human brain: high levels of cytochrome P4503A43 in brain and metabolism of anti-anxiety drug alprazolam to its active metabolite. PLoS One 3, e2337.

Ayano, G., 2016. Psychotropic Medications Metabolized by Cytochromes P450 (CYP) 2D6 Enzyme and Relevant Drug Interactions. Clinical Pharmacology & Biopharmaceutics 05.

Chen, Z.R., Irvine, R.J., Bochner, F., Somogyi, A.A., 1990. Morphine formation from codeine in rat brain: a possible mechanism of codeine analgesia. Life Sci 46, 1067–1074.

Cook, D., Brown, D., Alexander, R., March, R., Morgan, P., Satterthwaite, G., Pangalos, M.N., 2014. Lessons learned from the fate of AstraZeneca’s drug pipeline: a five-dimensional framework. Nat Rev Drug Discov 13, 419–431.

Decleves, X., Jacob, A., Yousif, S., Shawahna, R., Potin, S., Scherrmann, J.M., 2011. Interplay of drug metabolizing CYP450 enzymes and ABC transporters in the blood-brain barrier. Curr Drug Metab 12, 732–741.

Deepika, D., Kumar, V., 2023. The Role of “Physiologically Based Pharmacokinetic Model (PBPK)” New Approach Methodology (NAM) in Pharmaceuticals and Environmental Chemical Risk Assessment. International Journal of Environmental Research and Public Health 20.

Durairaj, P., Li, S., 2022. Functional expression and regulation of eukaryotic cytochrome P450 enzymes in surrogate microbial cell factories. Engineering Microbiology 2.

Fang, J., 2000. Metabolism of clozapine by rat brain: the role of flavin-containing monooxygenase (FMO) and cytochrome P450 enzymes. Eur J Drug Metab Pharmacokinet 25, 109–114.

Fanni, D., Pinna, F., Gerosa, C., Paribello, P., Carpiniello, B., Faa, G., Manchia, M., 2021. Anatomical distribution and expression of CYP in humans: Neuropharmacological implications. Drug Dev Res 82, 628–667.

Grobe, N., Kutchan, T.M., Zenk, M.H., 2012. Rat CYP2D2, not 2D1, is functionally conserved with human CYP2D6 in endogenous morphine formation. FEBS Lett 586, 1749–1753.

Hammarlund-Udenaes, M., Friden, M., Syvanen, S., Gupta, A., 2008. On the rate and extent of drug delivery to the brain. Pharm Res 25, 1737–1750.

Jolivalt, C., Minn, A., Vincent-Viry, M., Galteau, M.M., Siest, G., 1995. Dextromethorphan O-demethylase activity in rat brain microsomes. Neurosci Lett 187, 65–68.

Kadry, H., Noorani, B., Cucullo, L., 2020. A blood-brain barrier overview on structure, function, impairment, and biomarkers of integrity. Fluids Barriers CNS 17, 69.

Khokhar, J.Y., Tyndale, R.F., 2011. Drug metabolism within the brain changes drug response: selective manipulation of brain CYP2B alters propofol effects. Neuropsychopharmacology 36, 692–700.

Lipscomb, J.C., Poet, T.S., 2008. In vitro measurements of metabolism for application in pharmacokinetic modeling. Pharmacology & Therapeutics 118, 82–103.

Loryan, I., Reichel, A., Feng, B., Bundgaard, C., Shaffer, C., Kalvass, C., Bednarczyk, D., Morrison, D., Lesuisse, D., Hoppe, E., Terstappen, G.C., Fischer, H., Di, L., Colclough, N., Summerfield, S., Buckley, S.T., Maurer, T.S., Friden, M., 2022. Unbound Brain-to-Plasma Partition Coefficient, K(p,uu,brain)-a Game Changing Parameter for CNS Drug Discovery and Development. Pharm Res 39, 1321–1341.

Marsh, J.C., Chowdry, J., Parry-Jones, N., Ellis, S.W., Muir, K.R., Gordon-Smith, E.C., Tucker, G.T., 1999. Study of the association between cytochromes P450 2D6 and 2E1 genotypes and the risk of drug and chemical induced idiosyncratic aplastic anaemia. Br J Haematol 104, 266–270.

McMillan, D., 2018. Brain CYP2D Metabolism of Opioids Impacts Brain Levels, Analgesia, and Tolerance, Department of Pharmacology and Toxicology. University of Toronto.

McMillan, D.M., Tyndale, R.F., 2018. CYP-mediated drug metabolism in the brain impacts drug response. Pharmacol Ther 184, 189–200.

Miksys, S., Rao, Y., Sellers, E.M., Kwan, M., Mendis, D., Tyndale, R.F., 2000. Regional and cellular distribution of CYP2D subfamily members in rat brain. Xenobiotica 30, 547–564.

Miksys, S., Tyndale, R.F., 2013. Cytochrome P450-mediated drug metabolism in the brain. J Psychiatry Neurosci 38, 152–163.

Miksys, S.L., Tyndale, R.F., 2002. Drug-metabolizing cytochrome P450s in the brain. J Psychiatry Neurosci 27, 406–415.

Movin-Osswald, G., Boelaert, J., Hammarlund-Udenaes, M., Nilsson, L.B., 1993. The pharmacokinetics of remoxipride and metabolites in patients with various degrees of renal function. Br J Clin Pharmacol 35, 615–622.

Ogren, S.O., Lundstrom, J., Nilsson, L.B., 1993. Concentrations of remoxipride and its phenolic metabolites in rat brain and plasma. Relationship to extrapyramidal side effects and atypical antipsychotic profile. J Neural Transm Gen Sect 94, 199–216.

Ouzzine, M., Gulberti, S., Ramalanjaona, N., Magdalou, J., Fournel-Gigleux, S., 2014. The UDP-glucuronosyltransferases of the blood-brain barrier: their role in drug metabolism and detoxication. Front Cell Neurosci 8, 349.

R Core Team, 2023. R: A Language and Environment for Statistical Computing (4.3.0), 4.3.0 ed, p. R Foundation for Statistical Computing.

Sakakibara, Y., Katoh, M., Imai, K., Kondo, Y., Asai, Y., Ikushiro, S., Nadai, M., 2016. Expression of UGT1A subfamily in rat brain. Biopharm Drug Dispos 37, 314–319.

Saleh, M.A.A., Loo, C.F., Elassaiss-Schaap, J., De Lange, E.C.M., 2021. Lumbar cerebrospinal fluid-to-brain extracellular fluid surrogacy is context-specific: insights from LeiCNS-PK3.0 simulations. J Pharmacokinet Pharmacodyn 48, 725–741.

Sanchez-Dengra, B., Gonzalez-Alvarez, I., Bermejo, M., Gonzalez-Alvarez, M., 2021. Physiologically Based Pharmacokinetic (PBPK) Modeling for Predicting Brain Levels of Drug in Rat. Pharmaceutics 13.

Šrejber, M., Navrátilová, V., Paloncýová, M., Bazgier, V., Berka, K., Anzenbacher, P., Otyepka, M., 2018. Membrane-attached mammalian cytochromes P450: An overview of the membrane’s effects on structure, drug binding, and interactions with redox partners. Journal of Inorganic Biochemistry 183, 117–136.

Stevens, J., Ploeger, B.A., van der Graaf, P.H., Danhof, M., de Lange, E.C., 2011. Systemic and direct nose-to-brain transport pharmacokinetic model for remoxipride after intravenous and intranasal administration. Drug Metab Dispos 39, 2275–2282.

Summerfield, S.G., Zhang, Y., Liu, H., 2016. Examining the Uptake of Central Nervous System Drugs and Candidates across the Blood-Brain Barrier. J Pharmacol Exp Ther 358, 294–305.

Tyndale, R.F., Li, Y., Li, N.Y., Messina, E., Miksys, S., Sellers, E.M., 1999. Characterization of cytochrome P-450 2D1 activity in rat brain: high-affinity kinetics for dextromethorphan. Drug Metab Dispos 27, 924–930.

van den Brink, W.J., Elassaiss-Schaap, J., Gonzalez-Amoros, B., Harms, A.C., van der Graaf, P.H., Hankemeier, T., de Lange, E.C.M., 2017a. Multivariate pharmacokinetic/pharmacodynamic (PKPD) analysis with metabolomics shows multiple effects of remoxipride in rats. Eur J Pharm Sci 109, 431–440.

van den Brink, W.J., Wong, Y.C., Gulave, B., van der Graaf, P.H., de Lange, E.C., 2017b. Revealing the Neuroendocrine Response After Remoxipride Treatment Using Multi-Biomarker Discovery and Quantifying It by PK/PD Modeling. AAPS J 19, 274–285.

van der Lee, M., Allard, W.G., Vossen, R., Baak-Pablo, R.F., Menafra, R., Deiman, B., Deenen, M.J., Neven, P., Johansson, I., Gastaldello, S., Ingelman-Sundberg, M., Guchelaar, H.J., Swen, J.J., Anvar, S.Y., 2021. Toward predicting CYP2D6-mediated variable drug response from CYP2D6 gene sequencing data. Sci Transl Med 13.

Wang, Q., Han, X., Li, J., Gao, X., Wang, Y., Liu, M., Dong, G., Yue, J., 2015. Regulation of cerebral CYP2D alters tramadol metabolism in the brain: interactions of tramadol with propranolol and nicotine. Xenobiotica 45, 335–344.

Wang, Q., Zuo, Z., 2018. Impact of transporters and enzymes from blood-cerebrospinal fluid barrier and brain parenchyma on CNS drug uptake. Expert Opin Drug Metab Toxicol 14, 961–972.

Wang, W., Hallow, K.M., James, D.A., 2016. A Tutorial on RxODE: Simulating Differential Equation Pharmacometric Models in R. CPT Pharmacometrics Syst Pharmacol 5, 3–10.

Wishart, D.S., Feunang, Y.D., Guo, A.C., Lo, E.J., Marcu, A., Grant, J.R., Sajed, T., Johnson, D., Li, C., Sayeeda, Z., Assempour, N., Iynkkaran, I., Liu, Y., Maciejewski, A., Gale, N., Wilson, A., Chin, L., Cummings, R., Le, D., Pon, A., Knox, C., Wilson, M., 2018. DrugBank 5.0: a major update to the DrugBank database for 2018. Nucleic Acids Res 46, D1074–D1082.

Yamamoto, Y., Valitalo, P.A., Wong, Y.C., Huntjens, D.R., Proost, J.H., Vermeulen, A., Krauwinkel, W., Beukers, M.W., Kokki, H., Kokki, M., Danhof, M., van Hasselt, J.G.C., de Lange, E.C.M., 2018. Prediction of human CNS pharmacokinetics using a physiologically-based pharmacokinetic modeling approach. Eur J Pharm Sci 112, 168–179.

Yang, F., Liu, S., Wolber, G., Bureik, M., Parr, M.K., 2022. Complete Reaction Phenotyping of Propranolol and 4-Hydroxypropranolol with the 19 Enzymes of the Human UGT1 and UGT2 Families. Int J Mol Sci 23.

