## Supplementary material for "Exploring Kp,uu,BBB Values Smaller than Unity in Remoxipride: A Physiologically-Based CNS Model Approach Highlighting Brain Metabolism in Drugs with Passive Blood-Brain Barrier Transport": Table S1, Table S2, Figure S1, Figure S2, Table S3, Figure S3, Figure S4

**Journal:** European Journal of Pharmaceutical Sciences

**Mengxu Zhang^a^,** **Ilona M. Vuist^b^, Vivi Rottschäfer^c,d^, Elizabeth CM de Lange^a^**

a. Division of Systems Pharmacology and Pharmacy, Predictive Pharmacology group, Leiden Academic Centre of Drug Research, Leiden University, Leiden, The Netherlands
b. Charles River Laboratories, Groningen, The Netherlands
c. Mathematical Institute, Leiden University, Leiden, The Netherlands
d. Korteweg-de Vries Institute for Mathematics, University of Amsterdam, P.O. Box 94248, 1090 GE Amsterdam, The Netherlands

Correspondence :

Elizabeth CM de Lange,

Gorlaeus Laboratories, Einsteinweg 55, 2333 CC Leiden, The Netherlands

### 1. Experiments Details: Material and Methods for Experimental Assessment of Remoxipride

#### 1.1. Remoxipride

Remoxipride hydrochloride was obtained from Bio-Techne (Abington, United Kingdom). The molecular weight of remoxipride is 371.275 g/mol in the salt-free form, and 425.75 in the salt form, a correction factor of 1.15 was applied.

#### 1.2. *In vitro* experiments

*In vitro* experiments were performed to determine the recovery of remoxipride using MetaQuant microdialysis probes with a 3 mm exposed polyacrylonitrile membrane (MQ-PAN 3 mm, CRL Groningen, the Netherlands) and 3 mm exposed ethylene vinyl alcohol membrane (MQ-EC20 3 mm, CRL Groningen, the Netherlands). To this end, probes were placed in beakers containing 10 nM remoxipride in aCSF (147 mM NaCl, 3.0 mM KCl, 1.2 mM CaCl_2_, 1.2 mM MgCl_2_ in ultrapurified water). The beaker contents were continuously stirred and kept at a constant temperature of 37 °C. The probes were perfused with a slow flow of aCSF at a flow rate of 0.12 µL/min, and a carrier flow of ultrapurified H_2_O at a flow rate of 0.8 µL/min. After 2 hours of prestabilization, five consecutive microdialysis samples were collected in 15-minute intervals. Samples from the beaker content were collected at the start and end of the microdialysis experiment. All samples were collected into 300 µL polystyrene microvials (Microbiotech/se AB, Sweden; 4001029) and stored at -80 °C until analysis.

#### 1.3. *In vivo* experiments

##### 1.3.1.  Animals

A total of 13 adult male Sprague-Dawley rats (284 - 358 g; CRL Sulzfeld, Germany) were used for the experiments. The experiments were conducted in strict accordance with *the* Guide for the Care and Use of Laboratory Animals (National Research Council 2011) and were in accordance with European Union directive 2010/63 and the Dutch law. The experiments were carried out under license number AVD23600202011187, issued by the national committee for licensing of animal experiments (Centrale Commissie Dierproeven) and were approved by the Animal Care and Use Committee (Instantie voor Dierenwelzijn) of Charles River Discovery Groningen.

Following arrival, animals were housed in groups of up to 5 in polycarbonate cages (56 X 33 X 20 cm) with wire mesh top in a temperature (22 ± 2 °C) and humidity (55 ± 15%) controlled environment on a 12 hour light cycle (07.00 – 19.00h). After surgery, animals were housed individually (cages 30 X 30 X 30 cm). Standard diet (R/M-M A04 mod., Ssniff, Germany) and domestic quality mains water were available *ad libitum*.

##### 1.3.2.  Dose formulations

Remoxipride was formulated in saline at concentrations of 0.8 and 1.6 mg/mLfor i.v. administration of 4 and 8 mg/kg, in a volume of 5 mL/kg.

##### 1.3.3.  Experimental design

Rremoxipride was administered at t = 0 minutes. Experimental samples were collected 240 and 320 minutes after remoxipride administration. Samples were collected into polystyrene microvials (Microbiotech/se AB, Sweden; 4001029) using an automated fraction collector (UV 8301501, TSE, Univentor, Malta). During the experiment blood samples (50 µL) were collected via the jugular vein catheter according to the sampling schedule(0, 15, 30, 60, 90, 120, 180, 240, 300 min). All blood samples were collected into lithium heparin collection vials and plasma was collected following centrifugation into polystyrene vials. All samples were stored at ‑80°C. Terminal CSF was collected, weight was registered, and samples were stored at -80 °C. Histological verification of the probe position was not performed.

#### 1.4. Quantification of remoxipride

Concentrations of remoxipride were determined by HPLC with tandem mass spectrometry (MS/MS) detection. Plasma samples were mixed with acetonitrile, formic acid and internal standard (glafenine). Plasma samples were further precipitated and supernatant was mixed with formic acid in ultrapurified H_2_O.

An aliquot of each analytical sample was injected onto the HPLC column by an automated sample injector (Shimadzu, Japan). Chromatographic separation was performed using a Kinetex XB‑C18 column (50 x 2.1 mm, 2.6 µm; Phenomenex, USA) held at a temperature of 40 °C. The mobile phases consisted of A: 0.1% formic acid in ultrapurified H_2_O and B: 0.1% formic acid in acetonitrile. Elution proceeded using a linear gradient of phases A and B at a total flow rate of 0.5 mL/min.

The MS analyses were performed using an API 4000 MS/MS system consisting of an API 4000 MS/MS detector and a Turbo Ion Spray interface (both from Applied Biosystems, The Netherlands). The acquisitions were performed in positive ionization mode, with optimized settings for the analytes. The instrument was operated in multiple-reaction-monitoring (MRM) mode. MRM transitions for the analytes are 371 (Q1) and 243 (Q3) for remoxipride, and 373 (Q1) and 281 (Q3) for glafenine.

Suitable in-run calibration curves were fitted using weighted (1/x) regression, and the sample concentrations were determined using these calibration curves. Accuracy was verified by quality control samples after each sample series. Data were calibrated and quantified using the Analyst™ data system (Applied Biosystems).

**Table S1 Physio-chemical parameters of remoxipride**

| **drug** | **Molecular Weight** | **LogP** | **pKa** | **pKb** | **Km** |
| --- | --- | --- | --- | --- | --- |
| Remoxipride | 371.28 g/mol ^a^ | 2.94 ^a^ | 13.06 ^a^ | 8.4 ^a^ | 1738.28 ng/ml ^b^ |

a. Data comes from Drugbank ^1^
b. Estimated using study 1 to 4 datasets. Model and parameters estimated were shown in the Figure S2 and Table S3.

**Table S2 LeiCNS-PK physiological parameters values of rats**

| **Species** | | **Rat values** | |
| --- | --- | --- | --- |
| **Parameter** | | **Value**  **(range)** | **Reference** |
| **Volumes (mL)** | Total brain (V_tot_) | 1.8^[1]^ | ^2, 3^ |
|  | Brain extracellular fluid (V_ECF_) | 0.36^[2]^ | ^4^ |
|  | Brain intracellular fluid (V_ICF_) | 1.44^[3]^ | ^4^ |
|  | Brain cell lysosomes (V_LYS_) | 0.018^[4]^ | ^5^ |
|  | Brain microvasculature (V_MV_) | 0.054^[5]^ | ^6^ |
|  | Total cerebrospinal fluid (V_CSF_) | 0.28^[6]^  (0.155 – 0.4) | ^7-9^ |
|  | Lateral ventricles (V_LV_) | 0.0075^[7]^  (0.003 – 0.015) | ^8, 10-12^ |
|  | 3^rd^ & 4^th^ ventricles (V_TFV_) | 0.0075  (0.003 – 0.015) |  |
|  | Cisterna magna (V_CM_) | 0.017^[8]^ | ^13, 14^ |
|  | Subarachnoid space (V_SAS_) | 0.135^[9]^ | ^15^ |
| **Volume fractions** | Total brain phospholipid volume fraction (Vphb) | 0.05 | ^16^ |
|  | Endoplasmic reticulum volume fraction (fV_ER_) | 0.35^[10]^ | ^17^ |
|  | Endoplasmic reticulum membrane volume fraction (fV_ERM_) | 0.55^[11]^  (0.51-0.61) | ^18^ |
|  | Mitochondria volume fraction (fV_MT_) | 0.1^j^ | ^19^ |
|  | Mitochondria membranes volume fraction (fV_MTM_) | 0.3^k^  (0.21-0.39) | ^18^ |
| **Flows (mL min^-1^)** | Cerebral blood flow (Q_CBF_) | 2.87^[12]^ | ^6, 20^ |
|  | Brain ECF bulk flow (Q_ECF_) | 0.0002  (0.18E^-3^ - 0. 2E^-3^) | ^21-23^ |
|  | CSF flow (Q_CSF_) | 0.0022  (0.18E^-2^ - 0.22E^-2^) | ^8, 24^ |
| **Surface areas (cm^2^)** | Blood brain barrier (SA_BBB_) | 155  (150 - 188) | ^25-27^ |
|  | Blood CSF barrier (SA_BCSFB_) | 25^[13]^ | ^25^ |
|  | Brain cell plasma membrane (SA_BCM_) | 4250^[14]^ | ^4, 28^ |
|  | Lysosomes membrane (SA_LYS_) | 2700^[15]^ | ^29^ |
|  | Total brain cell phospholipid bilayer membrane (SA_BCphb_) | 212500^[16]^ | ^18^ |
| **Surface area fractions** | Endoplasmic reticulum surface area (rough and smooth) fraction (fSA_ER_) | 0.55^[17]^  (0.51-0.61) | ^18^ |
|  | Smooth endoplasmic reticulum surface area fraction (fSA_SER_) | 0.22^q^  (0.017-0.457) | ^18^ |
|  | Mitochondria outer membrane surface area fraction (fSA_MTout_) | 0.06^q^  (0.04-0.07) | ^18^ |
|  | Mitochondria inner membrane surface area fraction (fSA_MTinner_) | 0.2^q^  (0.17-0.32) | ^18^ |
| **Width**  **(µm)** | Blood brain barrier (W_BBB_) | 0.5  (0.2-0.5) | ^30^ |
|  | Blood CSF barrier (W_BCSFB_) |  |  |
| **Number** | Total brain cells (N_br,cells_) | 3.32E^8^ | ^28^ |
| **Pore size**  **(µm)** | Blood brain barrier (pTJ_BBB_) | 0.001 | ^31^ |
|  | Blood CSF barrier (pTJ_BCSFB_) | 0.009 | ^31^ |
| **Effective surface area  (%)** | BBB Transcellular transport (SA_BBB,T_) | 99.8^[18]^ | ^32-35^ |
|  | BCSFB Transcellular transport (SA_BCSFB,T_) | 99.8^n^ |  |
|  | BBB paracellular transport (SA_BBB,P_) | 0.006^[19]^ | ^31^ |
|  | BCSFB paracellular transport (SA_BCSFB,P_) | 0.05^o^ | ^31^ |
|  | Smooth endoplasmic reticulum metabolism fraction (efSA_SER_) | 0.013^[20]^ | ^36, 37^ |
|  | Mitochondria inner membrane metabolism fraction (efSA_MTinner_) | 0.2325^[21]^ | ^38, 39^ |
| **pH** | Plasma (pH_PL_) | 7.4 | ^40^ |
|  | Brain microvasculature (pH_MV_) |  |  |
|  | Brain extracellular fluid (pH_ECF_) | 7.3 | ^41^ |
|  | Cerebrospinal fluid (pH_CSF_) | 7.3 | ^42^ |
|  | Brain cells (pH_ICF_) | 7 | ^41^ |
|  | Brain cell lysosomes (pH_LYS_) | 5 | ^41^ |
|  | Endoplasmic reticulum (pH_ER_) | 7.2 | ^43^ |
|  | Mitochondria (pH_MT_) | 8 | ^43^ |
| **Metabolism (ng/ml*min)** | maximum velocity achieved by the brain enzyme when the drug concentration is high enough to saturate the enzyme (V_max_) | 39.26^[22]^ | estimated |

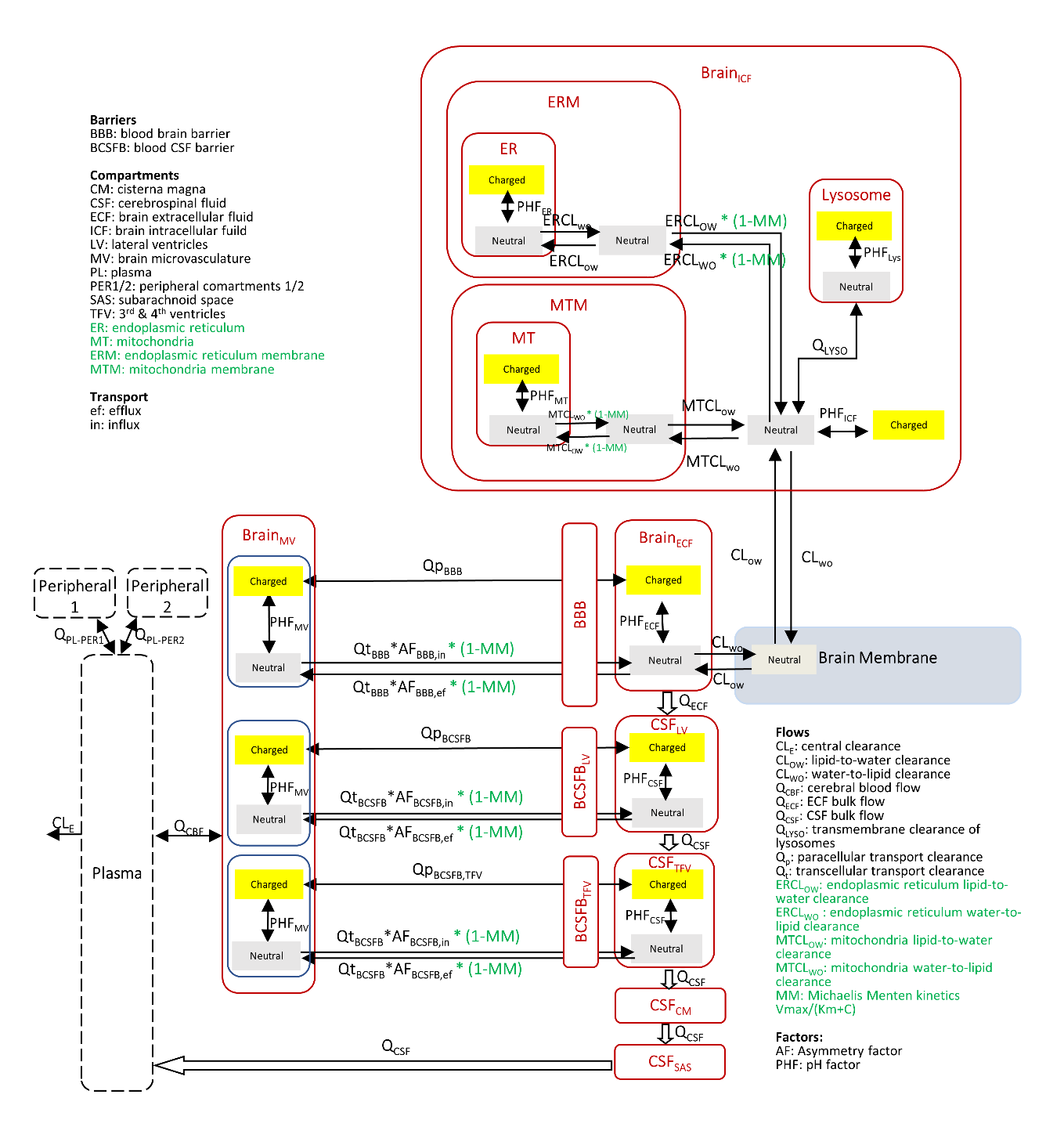

**Figure S1** **Detailed mathematical structure of LeiCNS-PK3.5.** *LeiCNS-PK3.0 is composed of whole body empirical plasma model and CNS PBPK model. Both models communicate via cerebral blood flow* ^44^. LeiCNS-PK3.5 *model incorporates brain metabolism into the flows through the BBB and BCSFB, and organelle’s level in brain ICF compartment. Green fonts represent the location of the metabolism incorporated.*

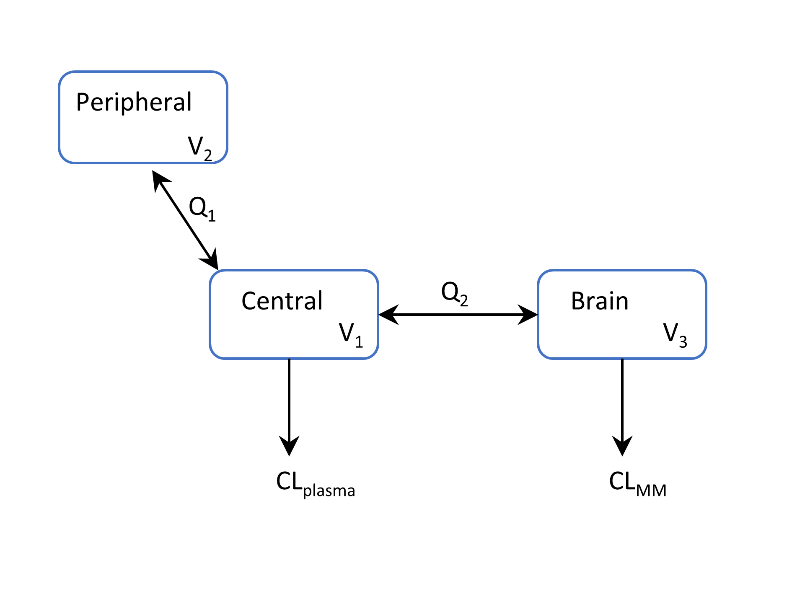

**Figure S2** **3-compartment model for estimating brain metabolism kinetic parameters.** *V_1_, volume of central compartment; V_2_, volume of peripherial compartment; V_3_, volume of brain compartment; Q_1_, flow between central and peripheral compartment; Q_2_, flow between central and brain compartment; CL_plasma_, clearance from central compartment. CL_MM_ represents the metabolism elimination from brain, which is consists of Km and Vmax using Michealis-Menten equation.*

**Table S3 Parameters of 3-compartment model for remoxipride brain metabolism.** *Parameters are corresponding to the Figure S2. In the central compartment,* *proportional residual variability was chosen to describe the differences between the observations and the individual predictions (Proportional_cen). In the brain compartment, proportional and additive residual variabilities were combined to describe differences between the observations and the individual predictions (Proportional_brain, Additional_brain). STOCH.APPROX, stochastic approximation; S.E., standard error; R.S.E, relative standard error. The parameters were estimated using the basic 3-compartmental model in Figure S2.*

| **REMOXIPRIDE** | **PARAMETERS** | **VALUES** | **STOCH.APPROX.** | |
| --- | --- | --- | --- | --- |
|  |  |  | **S.E.** | **R.S.E. (%)** |
| **Typical Value Estimates** | **CLplasma (mL/min)** | 7.55 | 0.83 | 11.0 |
|  | **Q1 (mL/min)** | 9.08 | 3.24 | 35.7 |
|  | **Q2 (mL/min)** | 35.44 | 2.67 | 7.53 |
|  | **V1 (mL)** | 326.42 | 17.85 | 5.47 |
|  | **V2 (mL)** | 36.72 | 2.17 | 5.92 |
|  | **V3 (mL)** | 3445.63 | 185.2 | 5.38 |
|  | **K_m_ (ng/ml)** | 39.26 | 1.47 | 3.76 |
|  | **V_max_ (ng/ml*min)** | 1738.28 | 108.11 | 6.22 |
| **Standard Deviation of the Interindividual Variability** | **Omega_CLplasma (mL/min)** | 0.64 | 0.088 | 13.7 |
|  | **Omega_Q1 (mL/min)** | 2.57 | 0.26 | 10.0 |
|  | **Omega_Q2 (mL/min)** | 0.5 | 0.068 | 13.7 |
|  | **Omega_V1 (mL)** | 0.33 | 0.048 | 14.3 |
|  | **Omega_V2 (mL)** | 0.52 | 0.048 | 9.21 |
|  | **Omega_V3 (mL)** | 0.46 | 0.041 | 8.93 |
|  | **Omega_K_m_ (ng/ml)** | 0.21 | 0.031 | 14.6 |
|  | **Omega_V_max_ (ng/ml*min)** | 0.54 | 0.049 | 9.03 |
| **Residual Variability** | **Proportional_cen** | 0.18 | 0.0073 | 3.96 |
|  | **Additional_brain** | 0.0001 | 0.0000078 | 7.85 |
|  | **Proportional_brain** | 0.26 | 0.0089 | 3.43 |

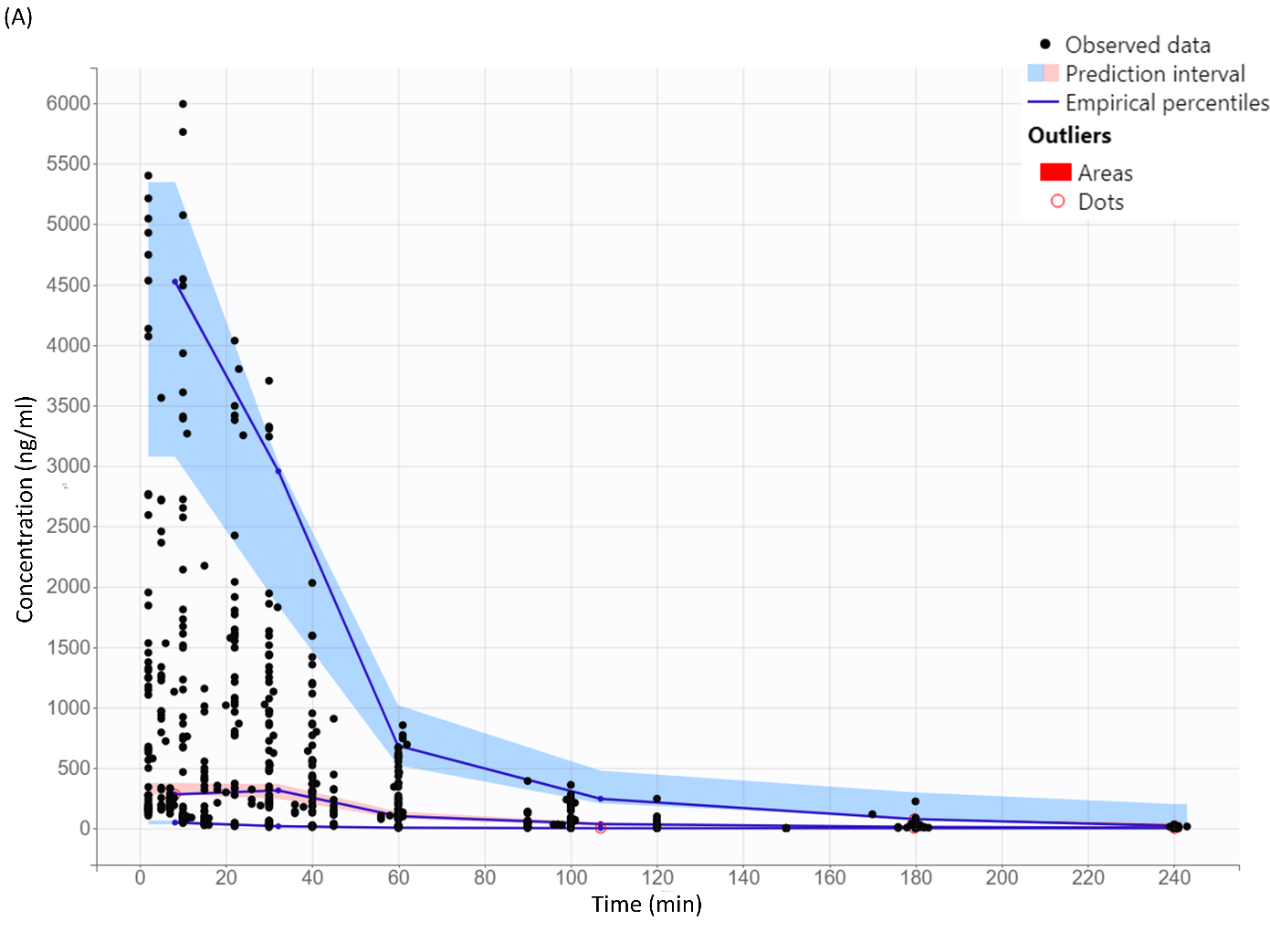

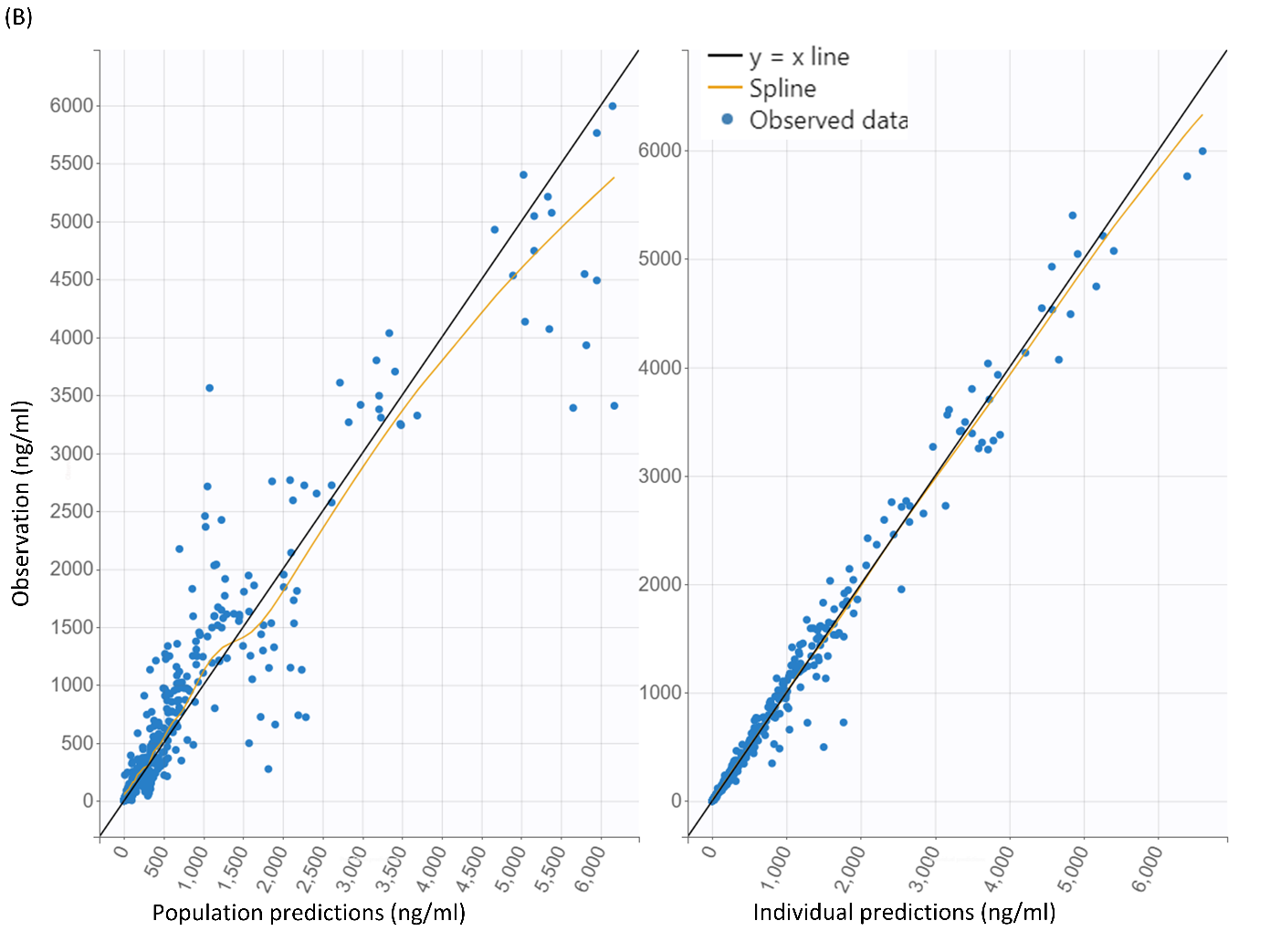

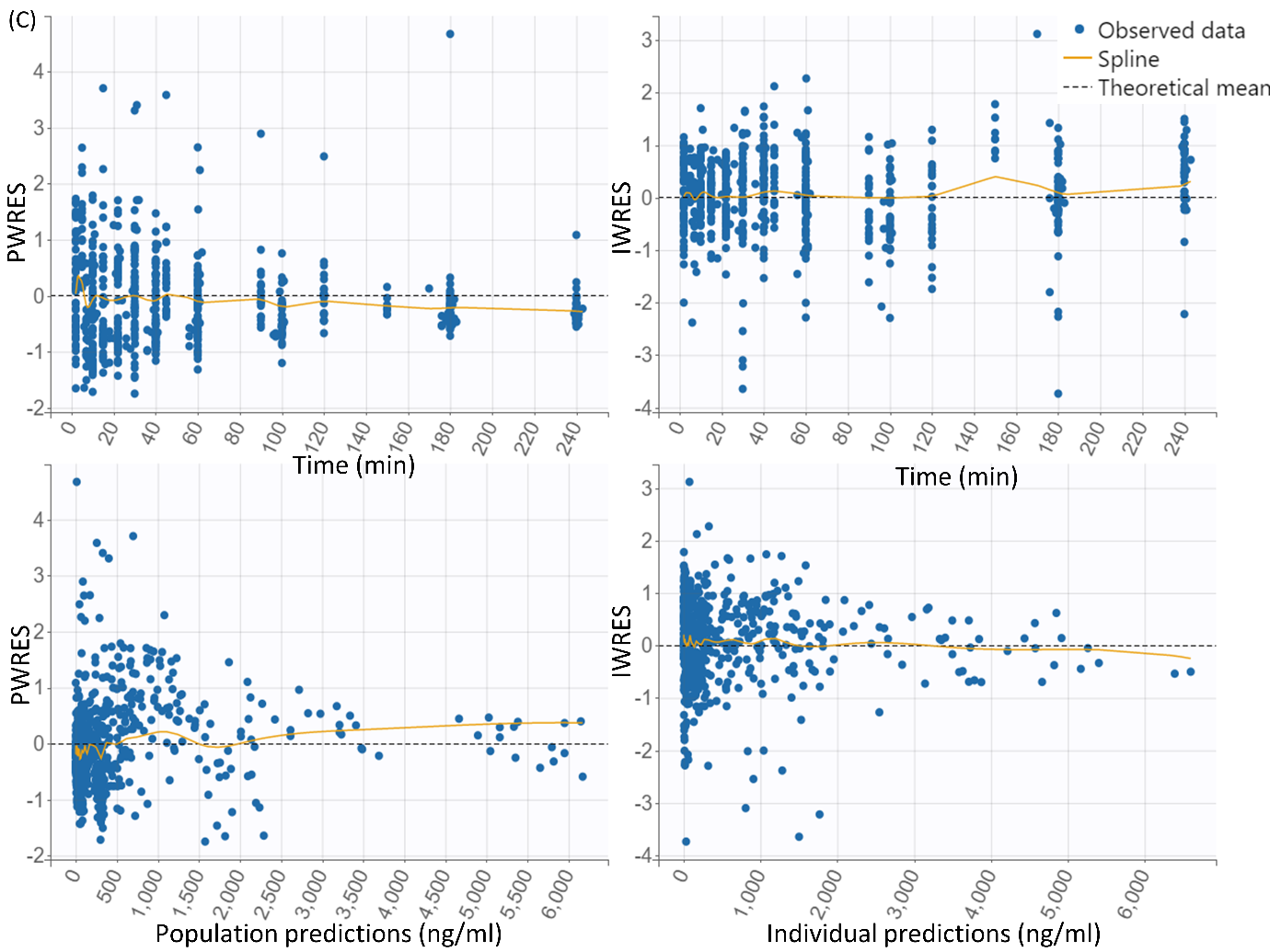

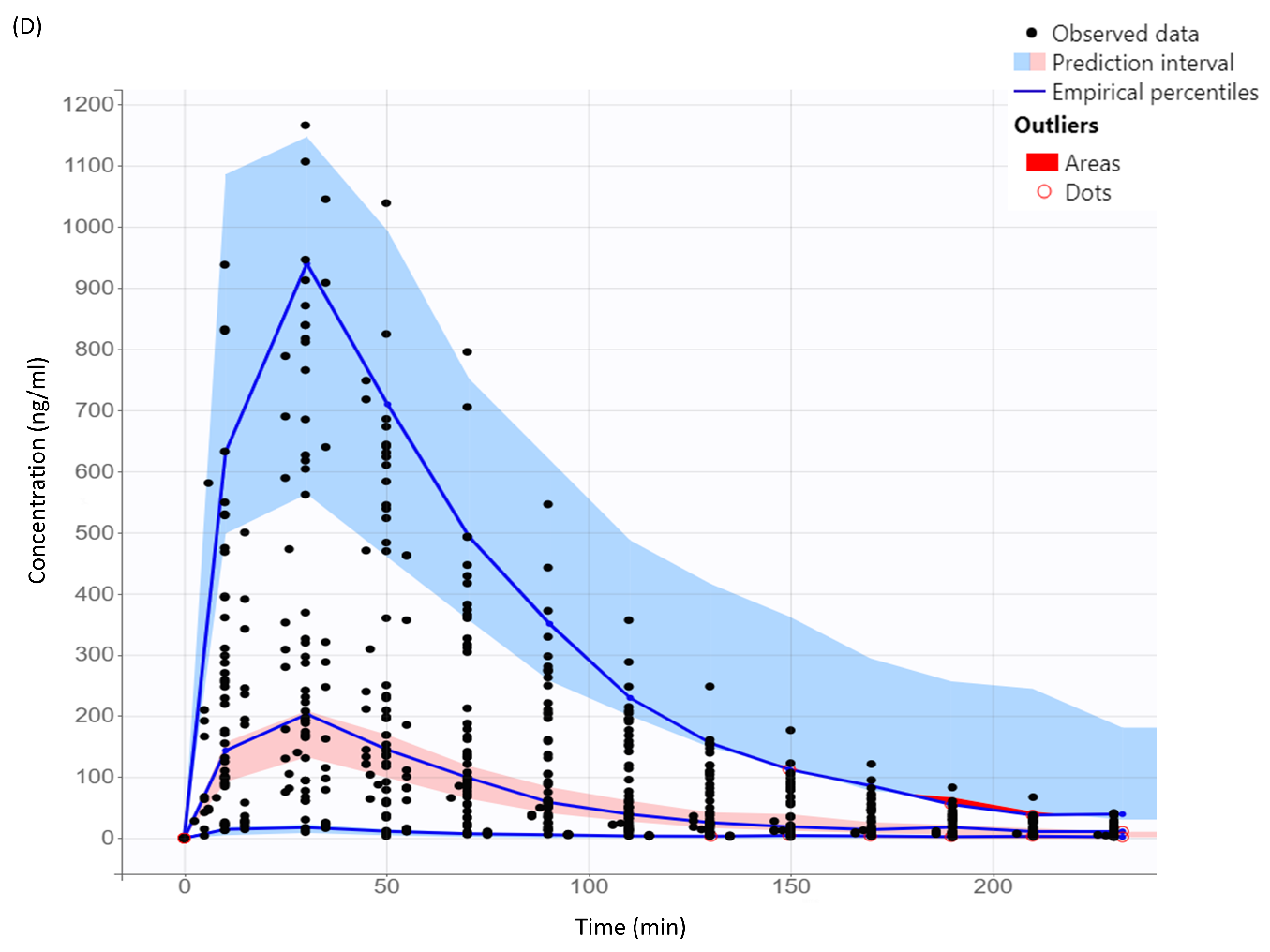

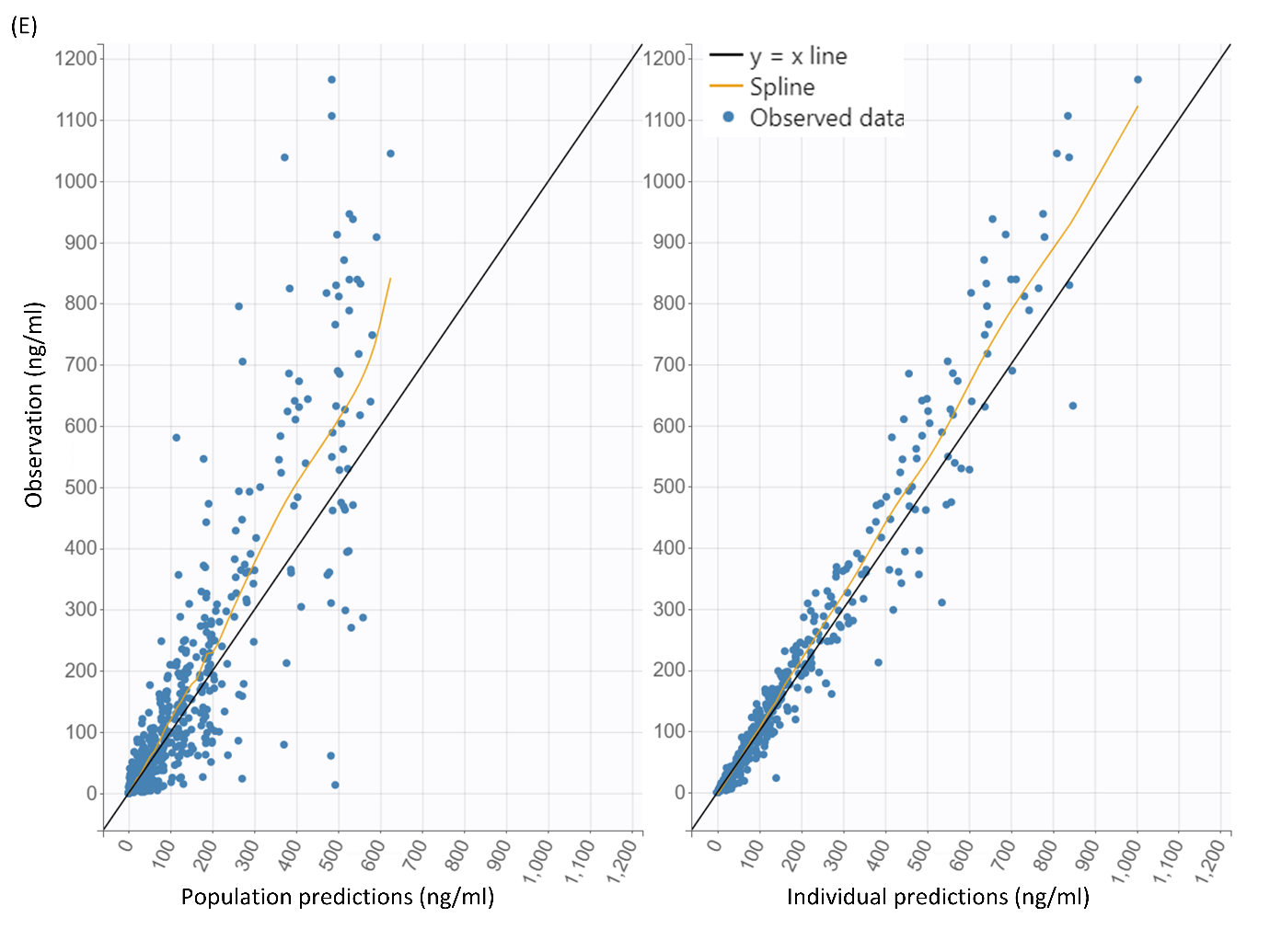

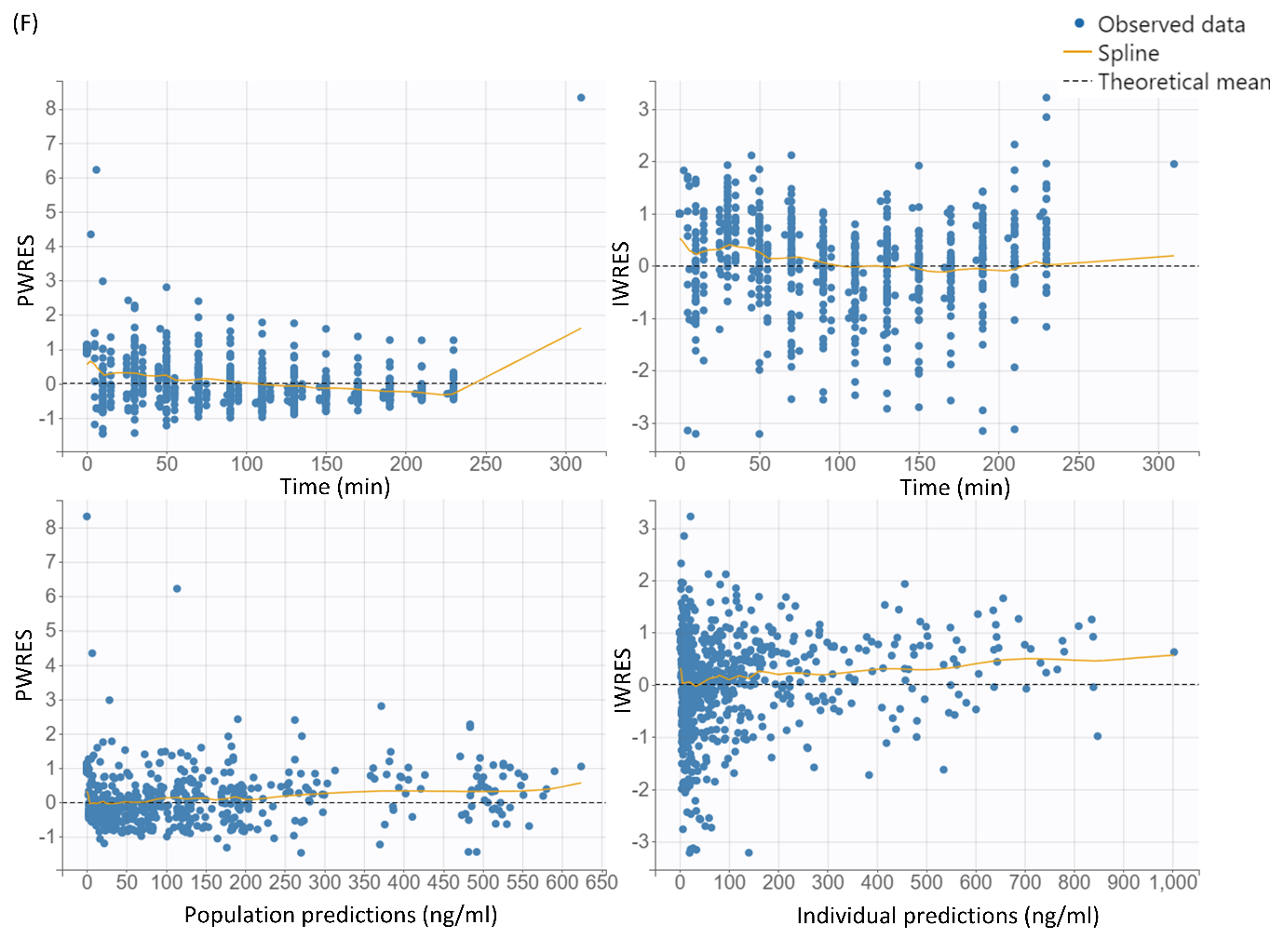

**Figure S3 Model evaluation plots of the 3-compartment brain metabolism model.** *A-C are evaluation figures for central compartment, D-F are evaluation figures for brain compartment. Visual predictive checks (VPCs) for the final model, based on 500 simulations in central (A) and brain compartment (D), which is corresponding to the Figure S2. The solid line connects the 10^th^, the 50^th^ and the 90^th^ percentiles of the observed values per bin from the up to the bottom. The blue areas represent the 95% confidence interval of the 10^th^ and 90^th^ percentiles. The pink area indicates the confidence interval of the 50^th^. The black points are the observation data points. The Y-axis is shown in a normal scale (A, D). Open circles and red areas represent the observed data are out of the 90% confidence interval of the simulated predictions. Diagnostic goodness-of-fit plots for the final model are also presented in (B, C) for central and (E, F) for brain compartment. (B, E) Observations versus population predictions; observations versus individual predictions; (C, F) population weighted residuals (PWRES) versus time after dose; PWRES versus population predictions; individual weighted residuals (IWRES) versus time after dose; IWRES versus individual predictions The locally weighted regression line (yellow spline), the line of unity (black solid lines) in B and E, and y = 0 (dashed lines) in C and F are shown.*

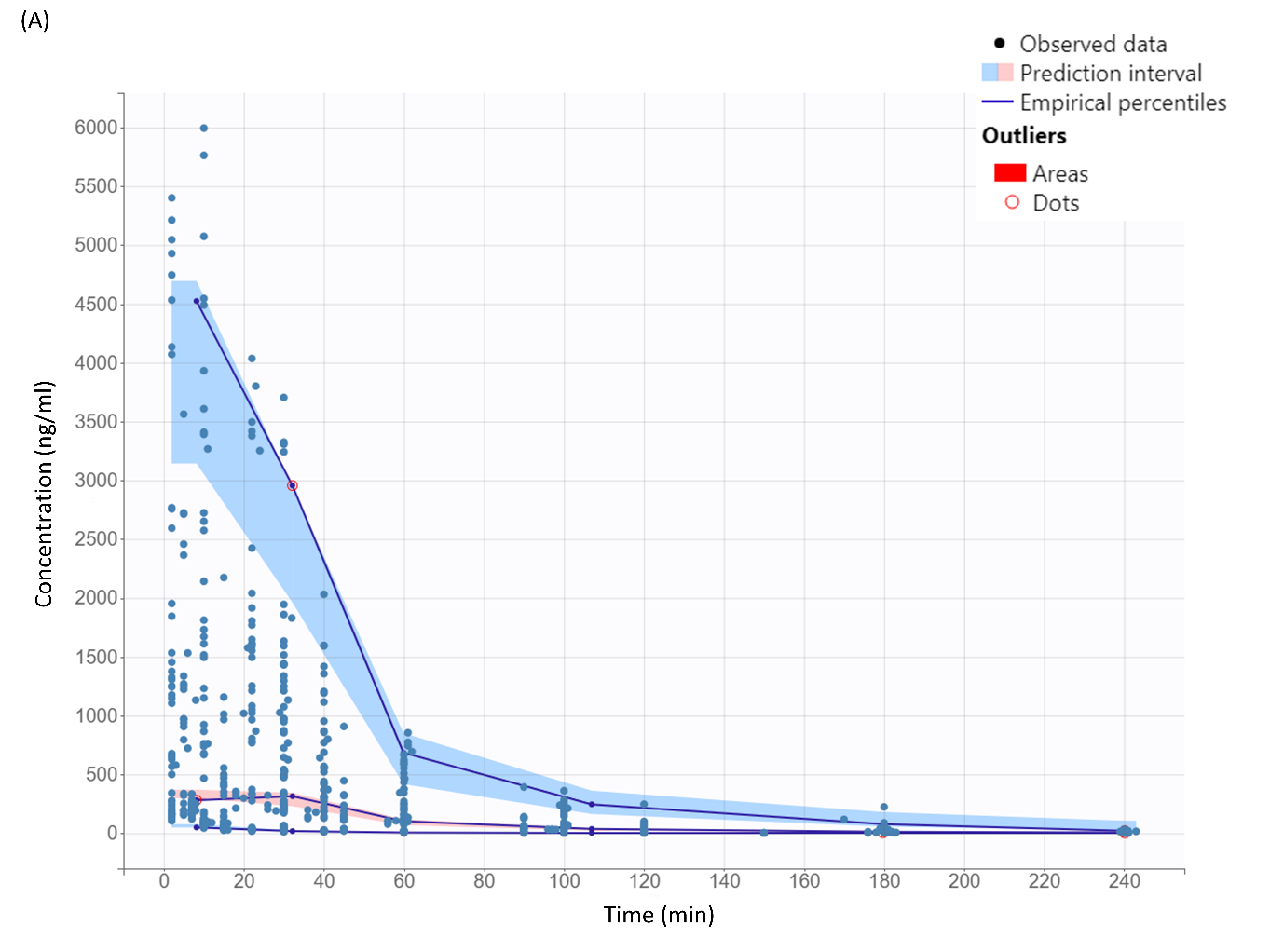

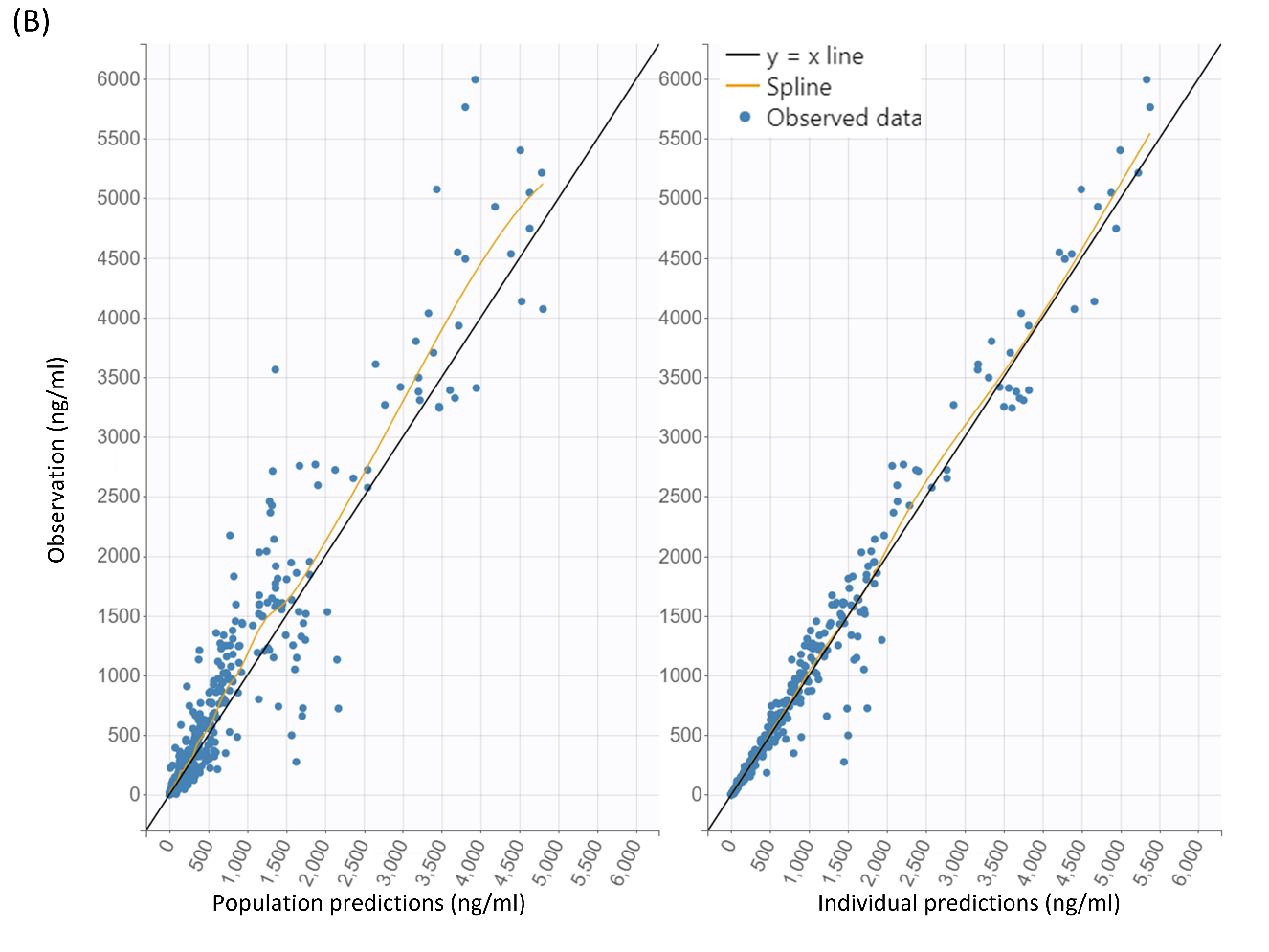

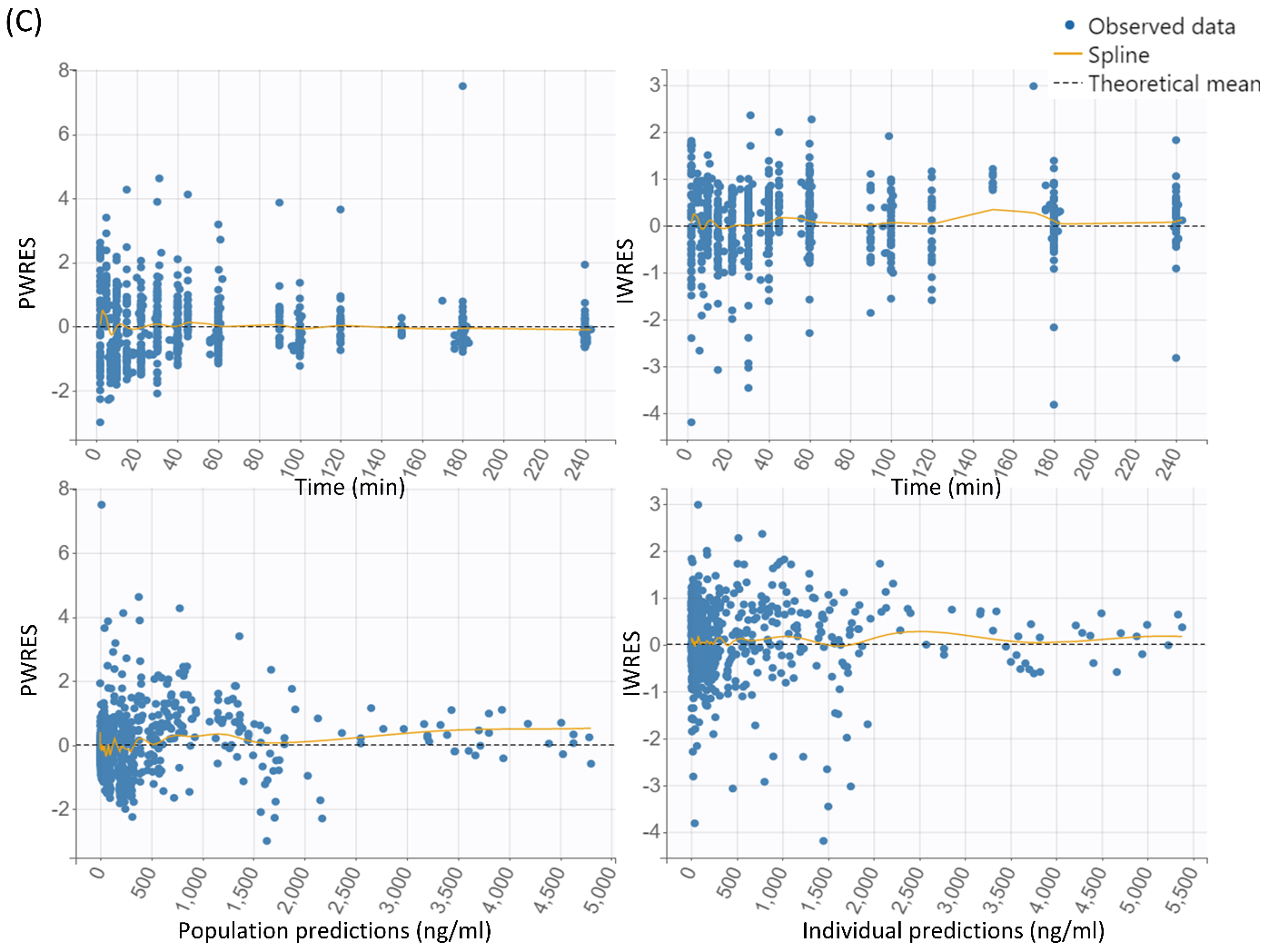

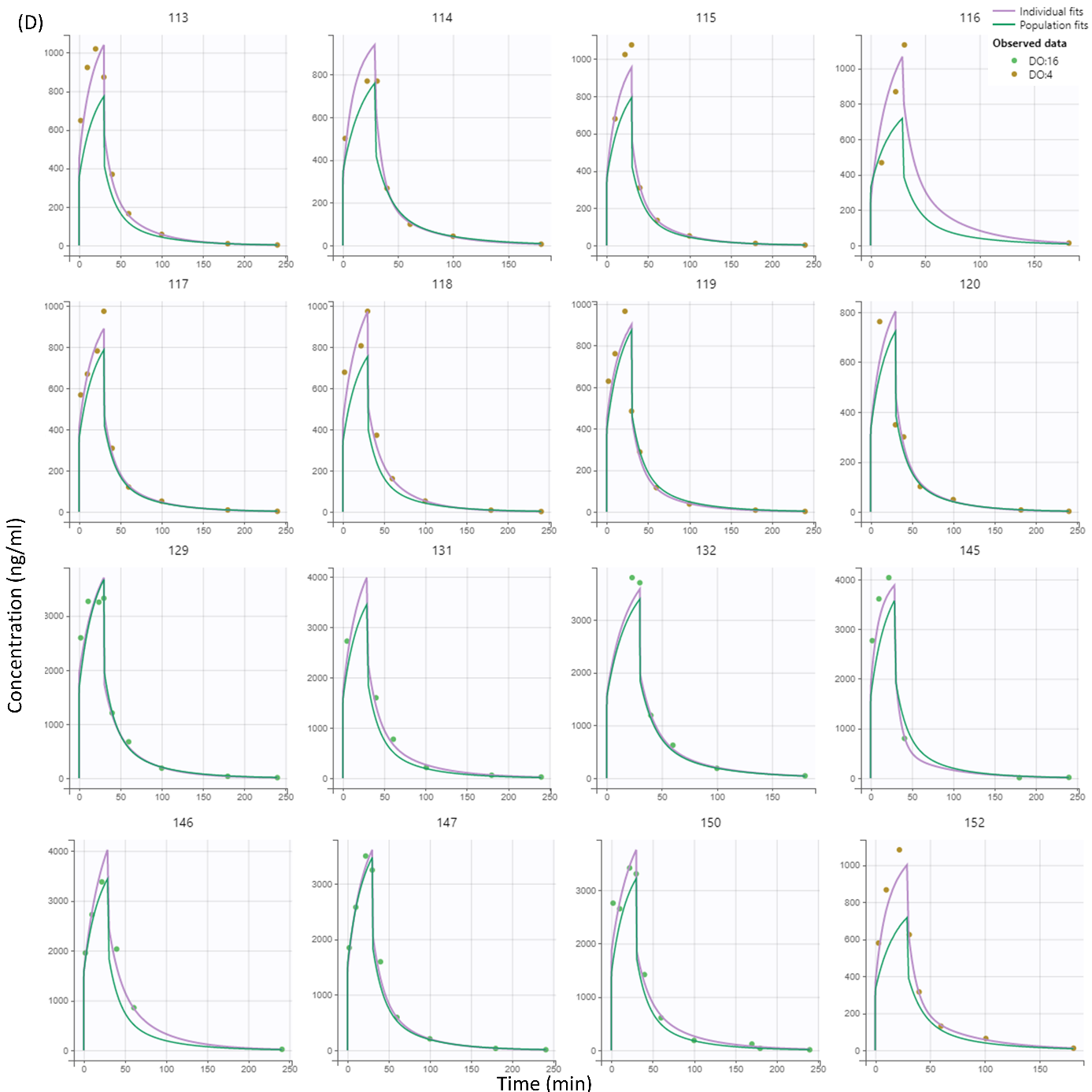

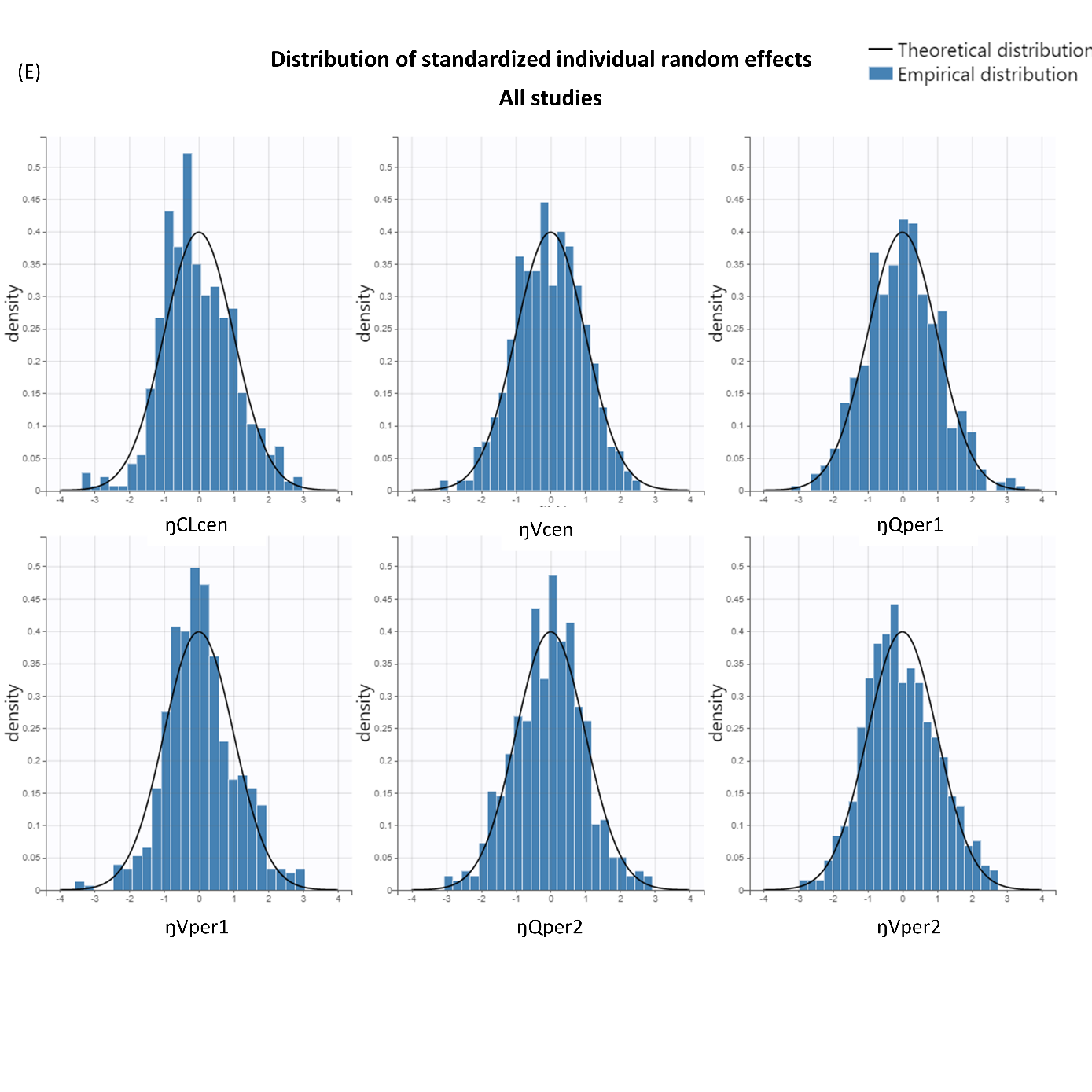

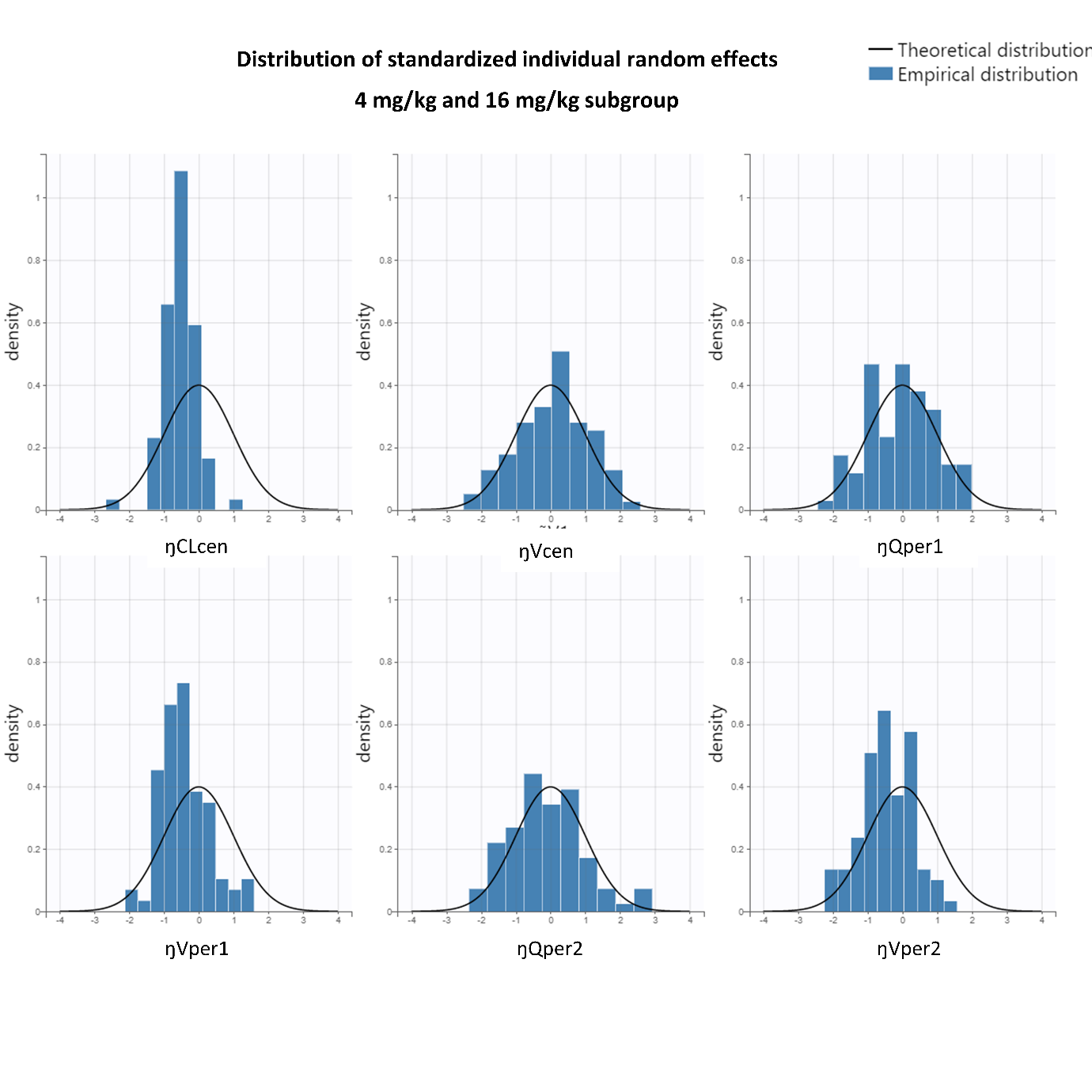

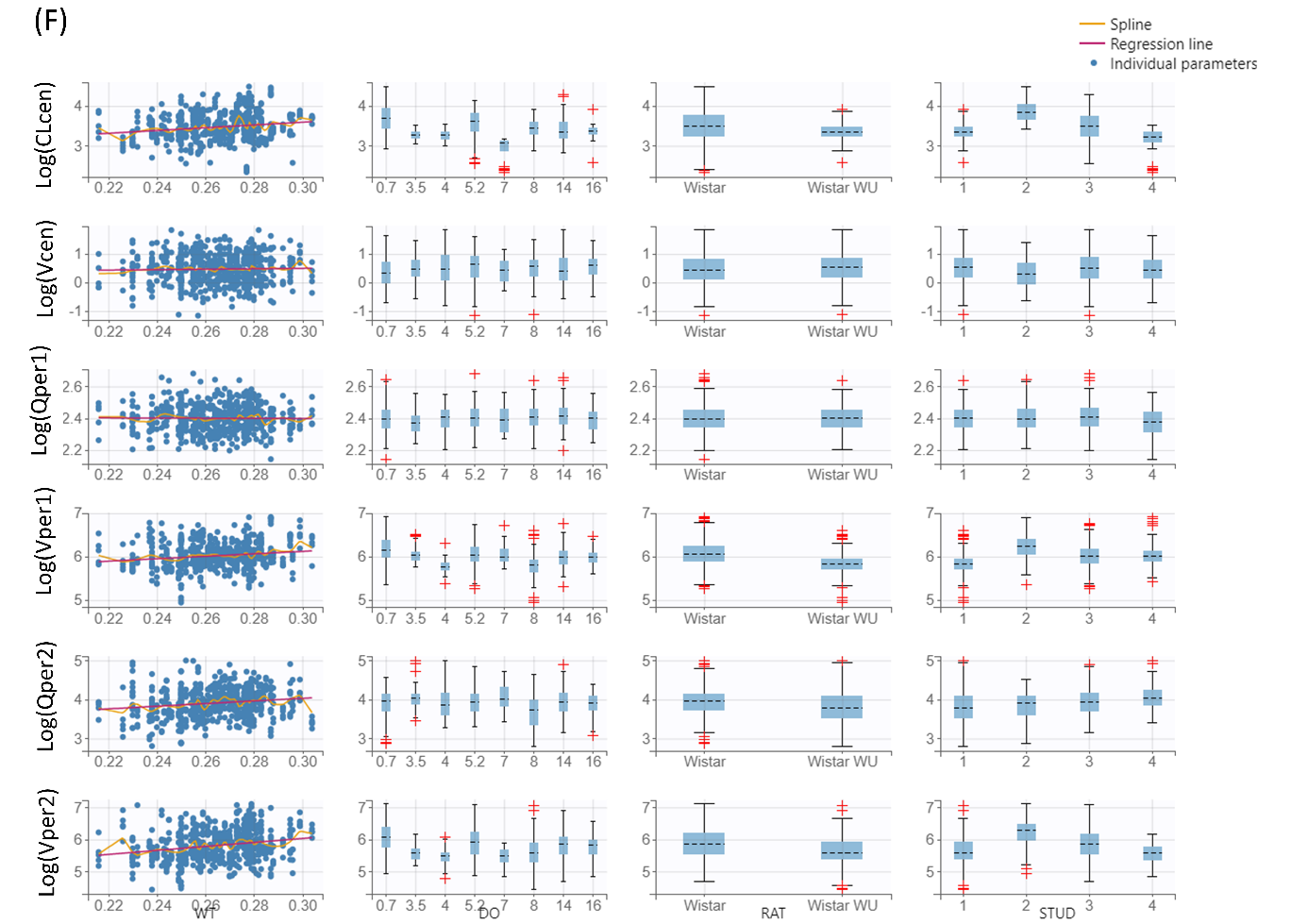

**Figure S4 Model evaluation plots of the rat plasma PK model.** *Visual predictive checks (VPCs) for the final model, based on 500 simulations in central compartment (A), which is corresponding to the Figure 3 . The solid line connects the 10^th^, the 50^th^ and the 90^th^ percentiles of the observed values per bin from the up to the bottom. The blue areas represent the 95% confidence interval of the 10^th^ and 90^th^ percentiles. The pink area indicates the confidence interval of the 50^th^. The black points are the observation data points. The Y-axis is shown in a normal scale (A). Open circles and red areas represent the observed data are out of the 90% confidence interval of the simulated predictions. Diagnostic goodness-of-fit plots for the final model are also presented in (B, C) for central compartment. (B) Observations versus population predictions; observations versus individual predictions; (C) population weighted residuals (PWRES) versus time after dose; PWRES versus population predictions; individual weighted residuals (IWRES) versus time after dose; IWRES versus individual predictions The locally weighted regression line (yellow spline), the line of unity (black solid lines) in B, and y = 0 (dashed lines) in C are shown. The individual concentration-time profiles’ fits are presented in (D) for each rat in study 1. Green solid lines present the population predictions, purple solid lines present individual predictions, yellow points are the observed data at dose 4 mg/kg; green points are the observed data at dose 8 mg/kg; and blue points are the observed data at dose 16 mg/kg. (E) is the distribution box plots for standardized individual random effects (ŋ) for all studies and the subgroup (4 mg/kg and 16 mg/kg). The individual parameters vs covariates plot (F) usually gives out the potential relevant covariates. However, in this plot, there is not a really significant relationship observed.*

**Supplementary references**

1.	Wishart DS, Feunang YD, Guo AC, et al. DrugBank 5.0: a major update to the DrugBank database for 2018. *Nucleic Acids Res*. Jan 4 2018;46(D1):D1074-D1082. doi:10.1093/nar/gkx1037

2.	Kawakami J, Yamamoto K, Sawada Y, Iga T. Prediction of brain delivery of ofloxacin, a new quinolone, in the human from animal data. *Journal of Pharmacokinetics and Biopharmaceutics*. 1994;22(3):207-227. doi:10.1007/bf02353329

3.	DiResta GR, Lee J, Arbit E. Measurement of brain tissue specific gravity using pycnometry. *Journal of Neuroscience Methods*. 1991;39(3):245-251. doi:10.1016/0165-0270(91)90103-7

4.	Lei Y, Han H, Yuan F, Javeed A, Zhao Y. The brain interstitial system: Anatomy, modeling, in vivo measurement, and applications. *Progress in Neurobiology*. 2017;157:230-246. doi:10.1016/j.pneurobio.2015.12.007

5.	Weibel ER, Stäubli W, Gnägi HR, Hess FA. Correlated Morphometric and Biochemical Studies on the Liver Cell. *The Journal of Cell Biology*. 1969;42(1):68-91. doi:10.1083/jcb.42.1.68

6.	Brown RP, Delp MD, Lindstedt SL, Rhomberg LR, Beliles RP. Physiological parameter values for physiologically based pharmacokinetic models. *Toxicology and Industrial Health*. Jul-Aug 1997;13(4):407-484. doi:Doi 10.1177/074823379701300401

7.	Preston JE. Ageing choroid plexus-cerebrospinal fluid system. *Microscopy Research and Technique*. 2001;52(1):31-37. doi:10.1002/1097-0029(20010101)52:1<31::Aid-jemt5>3.0.Co;2-t

8.	Chiu C, Miller MC, Caralopoulos IN, et al. Temporal course of cerebrospinal fluid dynamics and amyloid accumulation in the aging rat brain from three to thirty months. *Fluids Barriers CNS*. Jan 23 2012;9(1):3. doi:10.1186/2045-8118-9-3

9.	van den Berg MP, Romeijn SG, Verhoef JC, Merkus FW. Serial cerebrospinal fluid sampling in a rat model to study drug uptake from the nasal cavity. *J Neurosci Methods*. Apr 30 2002;116(1):99-107. doi:10.1016/s0165-0270(02)00033-x

10.	Basati S, Desai B, Alaraj A, Charbel F, Linninger A. Cerebrospinal fluid volume measurements in hydrocephalic rats. *J Neurosurg Pediatr*. Oct 2012;10(4):347-54. doi:10.3171/2012.6.PEDS11457

11.	Skjolding AD, Rowland IJ, Sogaard LV, Praetorius J, Penkowa M, Juhler M. Hydrocephalus induces dynamic spatiotemporal regulation of aquaporin-4 expression in the rat brain. *Cerebrospinal Fluid Res*. Nov 5 2010;7:20. doi:10.1186/1743-8454-7-20

12.	Levinger IM. The cerebral ventricles of the rat. *J Anat*. Apr 1971;108(Pt 3):447-51.

13.	Westerhout J, van den Berg DJ, Hartman R, Danhof M, de Lange EC. Prediction of methotrexate CNS distribution in different species - influence of disease conditions. *Eur J Pharm Sci*. Jun 16 2014;57:11-24. doi:10.1016/j.ejps.2013.12.020

14.	Lebedev SV, Blinov DV, Petrov SV. Spatial characteristics of cisterna magna in rats and novel technique for puncture with a stereotactic manipulator. *Bull Exp Biol Med*. Jun 2004;137(6):635-8. doi:10.1023/b:bebm.0000042732.00810.01

15.	Ghersi-Egea JF, Babikian A, Blondel S, Strazielle N. Changes in the cerebrospinal fluid circulatory system of the developing rat: quantitative volumetric analysis and effect on blood-CSF permeability interpretation. *Fluids Barriers CNS*. 2015;12:8. doi:10.1186/s12987-015-0001-2

16.	Poulin P, Theil FP. A Priori Prediction of Tissue:Plasma Partition Coefficients of Drugs to Facilitate the Use of Physiologically‐Based Pharmacokinetic Models in Drug Discovery. *Journal of Pharmaceutical Sciences*. 2000;89(1):16-35. doi:10.1002/(sici)1520-6017(200001)89:1<16::Aid-jps3>3.0.Co;2-e

17.	Valm AM, Cohen S, Legant WR, et al. Applying systems-level spectral imaging and analysis to reveal the organelle interactome. *Nature*. 2017;546(7656):162-167. doi:10.1038/nature22369

18.	Alberts B JA, Lewis J, et al. *Molecular Biology of the Cell. 4th edition. The Compartmentalization of Cells.* . New York: Garland Science; 2002.

19.	*Overflow Metabolism*. 2018.

20.	Sasaki Y, Wagner HN, Jr. Measurement of the distribution of cardiac output in unanesthetized rats. *J Appl Physiol*. Jun 1971;30(6):879-84. doi:10.1152/jappl.1971.30.6.879

21.	Szentistvanyi I, Patlak CS, Ellis RA, Cserr HF. Drainage of interstitial fluid from different regions of rat brain. *Am J Physiol*. Jun 1984;246(6 Pt 2):F835-44. doi:10.1152/ajprenal.1984.246.6.F835

22.	Mann JD, Butler AB, Johnson RN, Bass NH. Clearance of macromolecular and particulate substances from the cerebrospinal fluid system of the rat. *J Neurosurg*. Mar 1979;50(3):343-8. doi:10.3171/jns.1979.50.3.0343

23.	Abbott NJ. Evidence for bulk flow of brain interstitial fluid: significance for physiology and pathology. *Neurochem Int*. Sep 2004;45(4):545-52. doi:10.1016/j.neuint.2003.11.006

24.	Cserr H. Potassium exchange between cerebrospinal fluid, plasma, and brain. *Am J Physiol*. Dec 1965;209(6):1219-26. doi:10.1152/ajplegacy.1965.209.6.1219

25.	Keep RF, Jones HC. A morphometric study on the development of the lateral ventricle choroid plexus, choroid plexus capillaries and ventricular ependyma in the rat. *Brain Res Dev Brain Res*. Oct 1 1990;56(1):47-53. doi:10.1016/0165-3806(90)90163-s

26.	Sibbons PD, Aylward GL, Howard CV, vanVelzen D. A quantitative immunocytochemical analysis of total surface area of blood-brain barrier in developing rat brain. *Comparative Haematology International*. 1996;6(4):214-220. doi:Doi 10.1007/Bf00378113

27.	Hammarlund-Udenaes M, Friden M, Syvanen S, Gupta A. On the rate and extent of drug delivery to the brain. *Pharm Res*. Aug 2008;25(8):1737-50. doi:10.1007/s11095-007-9502-2

28.	Herculano-Houzel S, Mota B, Lent R. Cellular scaling rules for rodent brains. *Proc Natl Acad Sci U S A*. Aug 8 2006;103(32):12138-43. doi:10.1073/pnas.0604911103

29.	Bakker AC, Webster P, Jacob WA, Andrews NW. Homotypic fusion between aggregated lysosomes triggered by elevated [Ca2+]i in fibroblasts. *J Cell Sci*. Sep 1997;110 ( Pt 18):2227-38. doi:10.1242/jcs.110.18.2227

30.	Keep RF, Jones HC. Cortical microvessels during brain development: a morphometric study in the rat. *Microvasc Res*. Nov 1990;40(3):412-26. doi:10.1016/0026-2862(90)90036-q

31.	Sarin H. Physiologic upper limits of pore size of different blood capillary types and another perspective on the dual pore theory of microvascular permeability. *J Angiogenes Res*. Aug 11 2010;2:14. doi:10.1186/2040-2384-2-14

32.	Yamamoto Y, Valitalo PA, Huntjens DR, et al. Predicting Drug Concentration-Time Profiles in Multiple CNS Compartments Using a Comprehensive Physiologically-Based Pharmacokinetic Model. *CPT Pharmacometrics Syst Pharmacol*. Nov 2017;6(11):765-777. doi:10.1002/psp4.12250

33.	Haas TL, Duling BR. Morphology favors an endothelial cell pathway for longitudinal conduction within arterioles. *Microvasc Res*. Mar 1997;53(2):113-20. doi:10.1006/mvre.1996.1999

34.	Schulze C, Firth JA. Interendothelial junctions during blood-brain barrier development in the rat: morphological changes at the level of individual tight junctional contacts. *Brain Res Dev Brain Res*. Sep 18 1992;69(1):85-95. doi:10.1016/0165-3806(92)90125-g

35.	Weiss N, Miller F, Cazaubon S, Couraud PO. The blood-brain barrier in brain homeostasis and neurological diseases. *Biochim Biophys Acta*. Apr 2009;1788(4):842-57. doi:10.1016/j.bbamem.2008.10.022

36.	Smirle J, Au CE, Jain M, Dejgaard K, Nilsson T, Bergeron J. Cell biology of the endoplasmic reticulum and the Golgi apparatus through proteomics. *Cold Spring Harb Perspect Biol*. Jan 1 2013;5(1):a015073. doi:10.1101/cshperspect.a015073

37.	Fujisawa A, Tamura T, Yasueda Y, Kuwata K, Hamachi I. Chemical Profiling of the Endoplasmic Reticulum Proteome Using Designer Labeling Reagents. *J Am Chem Soc*. Dec 12 2018;140(49):17060-17070. doi:10.1021/jacs.8b08606

38.	Morgenstern M, Peikert CD, Lubbert P, et al. Quantitative high-confidence human mitochondrial proteome and its dynamics in cellular context. *Cell Metab*. Dec 7 2021;33(12):2464-2483 e18. doi:10.1016/j.cmet.2021.11.001

39.	Schenkel LC, Bakovic M. Formation and Regulation of Mitochondrial Membranes. *International Journal of Cell Biology*. 2014;2014:1-13. doi:10.1155/2014/709828

40.	Atherton JC. Acid-base balance: maintenance of plasma pH. *Anaesthesia & Intensive Care Medicine*. 2003;4(12):419-422. doi:10.1383/anes.4.12.419.27385

41.	Friden M, Bergstrom F, Wan H, et al. Measurement of unbound drug exposure in brain: modeling of pH partitioning explains diverging results between the brain slice and brain homogenate methods. *Drug Metab Dispos*. Mar 2011;39(3):353-62. doi:10.1124/dmd.110.035998

42.	Siesjo BK. Symposium on acid-base homeostasis. The regulation of cerebrospinal fluid pH. *Kidney Int*. May 1972;1(5):360-74. doi:10.1038/ki.1972.47

43.	Casey JR, Grinstein S, Orlowski J. Sensors and regulators of intracellular pH. *Nature Reviews Molecular Cell Biology*. 2009;11(1):50-61. doi:10.1038/nrm2820

44.	Saleh MAA, Loo CF, Elassaiss-Schaap J, De Lange ECM. Lumbar cerebrospinal fluid-to-brain extracellular fluid surrogacy is context-specific: insights from LeiCNS-PK3.0 simulations. *J Pharmacokinet Pharmacodyn*. Oct 2021;48(5):725-741. doi:10.1007/s10928-021-09768-7

1. Based on rat brain weight (1.88 gm) and density (1.04-1.05 gm ml**^-1^**)
2. Calculated as 15-20 (20 was used)% of total brain volume
3. Calculated as 80% of total brain volume
4. Calculated as 1.25% (1/80) of ICF volume; based on liver lysosomes
5. Calculated as 3% of total brain volume
6. Mean of the 4 values
7. Assuming equal volumes of the ventricles; based on volumes of three-month-old rats
8. [Calculated as 5.7% of total CSF volume and according to cisterna magna geometry](https://www.ncbi.nlm.nih.gov/pmc/articles/PMC4365764/)
9. Calculated as 48% of total CSF volume, based on measurement performed in 9-day-old rats
10. Fraction of ICF volume, based cortical neurons from eukaryotic animal.
11. Fraction of total brain phospholipid, based on total brain volume and V_phb_ , data is for the general eukaryotic animal cells
12. Calculated as 2.6% of total cardiac output
13. Based on three-month-old rats, surface area at lateral ventricles (and 3^rd^ and 4^th^ ventricles) is assumed 50% of total surface area
14. Based on ICF total volume, total number of brain cells, and assuming spherical cells to calculate the radius which is used with total number of brain cells to calculate total surface area of brain cell membranes
15. Based on lysosomes total volume and the average radius of rat kidney lysosomes (0.2 µm)
16. Calculated as 50 times of brain cell plasma membrane, based on eukaryotic animal cells
17. Fraction of total brain cell phospholipid bilayer membrane, based on eukaryotic animal cells
18. Based on relative length of intercellular space (0.03 μm) and cell perimeter (17 μm).
19. Based on an endothelial cell perimeter of 17 um

    ^P^ Based on the model estimation showed in Figure S1 and Table S3.
20. Based on surface area of smooth ER, protein fraction of smooth ER (43.1%) from Hela cells, and rat ER metabolism protein fraction (3%).
21. Based on surface area of inner membrane of mitochondria, protein percentage of mitochondria inner membrane (75%), and metabolism protein abundance (31%).
22. 3-compartment PopPK model estimated parameters from rat datasets, more details see Figure S1, Table S3 and Figure S3.
